# Integrative AlphaFold Modeling, Fragment Mapping, and Microsecond Molecular Dynamics Reveal Ligand-Specific Structural Plasticity at the Human Urotensin II Receptor

**DOI:** 10.64898/2026.04.04.716502

**Authors:** Alexandre G. Torbey

**Affiliations:** Institut National de la Recherche Scientifique (INRS), Centre Armand-Frappier Santé Biotechnologie, Laval, H7V 1B7, Canada

**Keywords:** G protein coupled receptor (GPCR), Urotensin II Receptor (UT), Cyclic Peptide, Structure-Activity Relationships, Site Identification by Ligand Competitive Saturation (SILCS), Molecular Dynamics Simulations, Peptide Ligands, Conformational Plasticity

## Abstract

Peptide ligands Urotensin II (*h*UII, human), *h*UII-related peptide (URP) and its cognate human receptor (*h*UT) are known for their implications in cardiovascular pathophysiology, yet the lack of experimentally resolved *h*UT structures has limited a deep mechanistic understanding of ligand binding and receptor activation. Here, we leverage recent breakthroughs in multistate AlphaFold predictions, long-timescale molecular dynamics (MD) simulations, and site identification by ligand competitive saturation (SILCS) based pocket mapping and solving ligand bound conformation to illuminate the dynamic interaction of *h*UII and URP with *h*UTR. By analyzing *h*UT dynamics in its intracellular transducer binding pocket, and residue-level interaction probabilities in each simulation, we capture subtle distinctions in the way *h*UII and URP anchor key pocket residues, modulate transmembrane (TM) domain tilts. Results indicate that *h*UII imposes stronger conformational constraints on TM5 and TM6 relative to URP, both potentially stabilizing different active-like receptor configurations. At the same time, interaction maps highlight unique aromatic and polar networks that each ligand exploits. These findings reinforce the concept that relatively small differences in GPCR peptide ligand structure may lead to large effects on receptor-state selection, signal specificity, ultimately reflecting different clinical outcomes. By integrating computational modeling with per-residue dynamics, this work not only reconciles prior mutagenesis and docking data but also provides validated 3D models and MD simulations of the endogenous ligands bound to *h*UT, offering new opportunities to selectively harness ligand-dependent signaling in the urotensinergic system.

## 1. Introduction

Human Urotensin II (*h*UII) and the structurally related urotensin II-related peptide (URP), both endogenous peptides of the urotensinergic system (sequences shown in **Figure_1**), have drawn considerable attention for their roles in regulating the cardiovascular system, through their interaction with the human urotensin II receptor (*h*UT), a class A G protein-coupled receptor (GPCR). Despite conflicting evidence regarding the therapeutic interest of targeting this system to alleviate cardiac and vascular maladaptive outcomes (such as hypertrophy and fibrosis), a robust body of evidence highlights the need to dissect the underlying receptor-ligand mechanisms to design fine-tuned derivatives with translational potential (Avagimyan et al., 2022; Esposito et al., 2011; Mazarura et al., 2022; Nassour et al., 2019; Pereira-Castro et al., 2019; Russell, 2008). Attention has focused on the phenomenon of “functional selectivity”, also known as “biased signaling”, the concept that distinct ligands can select or stabilize specific receptor conformations, thereby promoting the activation or deactivation of certain signaling pathways over others (e.g., β-arrestin over G protein-dependent signaling), with major therapeutic implications.

**Figure 1.**
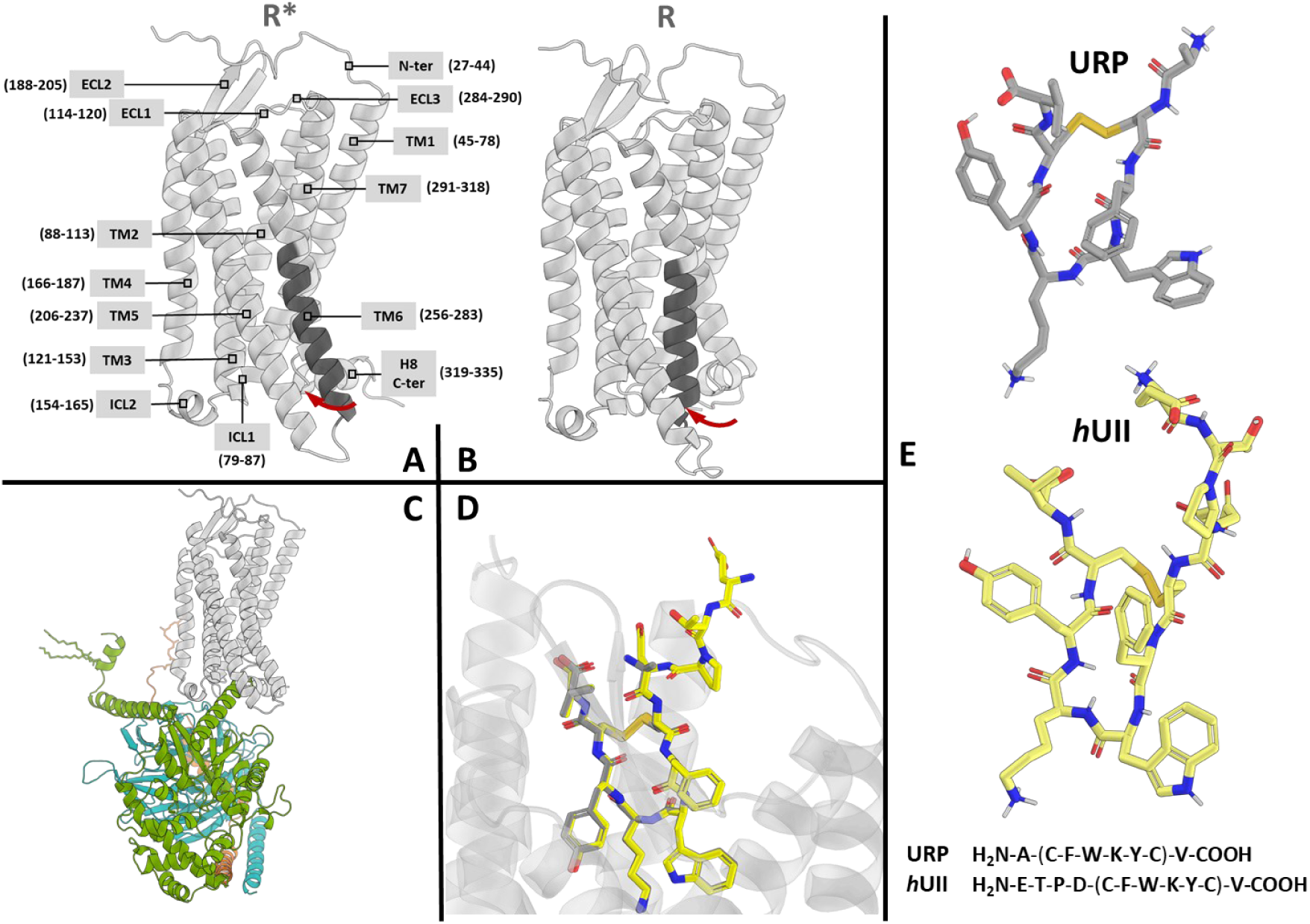
The urotensinergic system. Homology models of *h*UT in its inactive (R, A) and active (R*, B) conformations. The cytosolic end of TM6 is shown in dark grey, and its inward tilt is indicated by a red arrow. Residue boundaries for TM segments, loops and termini are provided in parentheses. The modeled *h*UT R*-G_q_ heterotetramer (C) includes human Gα_q_ (green), Gβ_2_ (blue) and Gγ_1_ (orange), with *h*UT shown in light grey. AlphaFold3 predictions of *h*UT bound to *h*UII (yellow) and URP (dark grey) (D) display nearly identical binding poses, indicating the model’s limited ability to distinguish between the two ligands. Panel E presents the sequences and three-dimensional structures of *h*UII and URP, with the disulfide-linked cyclic motifs indicated in parentheses; polar hydrogens are shown in white.

Several in vitro studies illustrate this concept. Recent advances in GPCR pharmacology highlight biased agonism as a promising approach to selectively modulate intracellular signalling. Clinically, carvedilol exemplifies this principle by inhibiting G protein-dependent cascades while engaging β-arrestin pathways at β-adrenergic receptors, an effect associated with enhanced cardioprotection and reduced apoptosis in heart failure models (Wisler et al., 2007). Likewise, oliceridine (TRV130), a μ-opioid receptor agonist, preferentially activates G-protein signalling over β-arrestin recruitment, resulting in effective analgesia with diminished respiratory depression and gastrointestinal side effects (Gan & Wase, 2020; Lambert & Calo, 2020). These examples demonstrate how pathway-selective receptor activation (e.g., via biased agonism) can improve therapeutic precision and minimize adverse events, thereby expanding GPCR-based drug discovery. These findings underscore how a single receptor can engage distinct downstream pathways, potentially affecting clinical outcomes.

Beyond classical potency and efficacy considerations, *h*UII and URP display clear examples of endogenous functional selectivity at *h*UT, supporting the view that closely related peptides can drive divergent biological responses. Several early pharmacological studies demonstrated ligand-specific differences across multiple physiological systems. In rat heart nuclei, *h*UII and URP differentially regulate transcription initiation, indicating distinct nuclear signalling capacities; similarly, only *h*UII stimulates polyphosphoinositide turnover and astrocyte proliferation in primary rat astrocytes, whereas URP elicits weaker or qualitatively different responses in the same preparations (Jarry et al., 2010; Johns et al., 2004). Functional divergence also emerges in cardiovascular models: URP is more potent than *h*UII in triggering Atrial Natriuretic Peptide release and modulating myocardial contractility in isolated rat heart preparations (Nassour et al., 2019), whereas *h*UII more strongly engages pathways linked to metabolic remodeling. In human aortic valve disease, the two peptides also appear biologically dissociated: URP mRNA is selectively upregulated in calcified fibrotic valves, and URP immunostaining correlates with regions of active calcification, while *h*UII shows no such enrichment (Nassour et al., 2019). Conversely, *h*UII, but not URP, reduces APOA1-dependent cholesterol efflux in M2 macrophages, downregulates ABCA1 in human aortic valve interstitial cells, enhances β-catenin nuclear accumulation, and promotes mineralization of these cells (Liu et al., 2009; Shi et al., 2016; Tzanidis et al., 2003). Finally, in acute heart failure patients, circulating URP levels greatly exceed those of *h*UII, suggesting differential regulation of both peptides in pathological states. Altogether, these findings underscore that endogenous *h*UT ligands are inherently biased, shaping distinct transcriptional, metabolic, and tissue-level outcomes despite their closely related structures.

However, how such in vitro differences translate into distinct molecular interactions between *h*UT and its cognate ligands is unknown. Harnessing such functional selectivity could enhance the pharmacological success of therapies targeting *h*UT in cardiovascular disease through the rational design of β-arrestin-biased ligands. For instance, Esposito *et al*. (2011) demonstrated that *h*UII-induced transactivation of the epidermal growth factor receptor (EGFR), mediated by β-arrestin 1/2, can confer a cardioprotective effect under pressure-overload stress. Accumulating evidence further supports the therapeutic interest of β-arrestin-biased *h*UT ligands, particularly in the cardiovascular context. Multiple studies have linked the deleterious hypertrophic and profibrotic actions of *h*UT activation to Gq-dependent signalling, mirroring the pathological Gq-driven responses described for the angiotensin AT1 receptor and β-adrenergic receptors (Liu et al., 2009; Shi et al., 2016; Tzanidis et al., 2003). In contrast, β-arrestin-mediated pathways appear protective: *h*UII-induced EGFR transactivation, which operates via β-arrestin 1/2 scaffolding, promotes cardiomyocyte survival and mitigates pressure-overload-induced dysfunction (Esposito et al., 2011). Consistent with this protective role, an analogue unable to recruit β-arrestin, urantide, fails to trigger EGFR transactivation and instead exacerbates cardiac dysfunction under mechanical stress (Esposito et al., 2011). Moreover, selective inhibition of GRK5, a kinase that preferentially phosphorylates *h*UT to promote Gq-dependent desensitization and pathological hypertrophic signalling, attenuates *h*UII-induced hypertrophy in neonatal cardiomyocytes (Park et al., 2016). These observations collectively support the therapeutic concept that *h*UT ligands able to avoid Gq/GRK5-dominated pathways while promoting β-arrestin recruitment and EGFR transactivation may offer improved cardioprotective profiles, mirroring biased-agonism strategies already in clinical use for other GPCRs. Beyond physiology, structure-activity relationships (SAR) studies have provided insight into key pharmacophore structural features of the *h*UII and URP peptides. Substitutions either in the cyclic region (Cys-Phe-Trp-Lys-Tyr-Cys) or in the flexible N-terminal extension have revealed how minor sequence alterations can markedly influence potency, efficacy, and signaling outcomes (É. Billard et al., 2019; É. Billard & Chatenet, 2020; Brancaccio et al., 2015; Chatenet et al., 2004, 2006; Merlino et al., 2019; Nassour et al., 2019).

Yet, *in silico* studies of these two endogenous ligands have historically been hampered by the lack of a high-resolution, experimentally determined *h*UT structure. The field has relied on homology models and docking approaches which, while informative, are limited in their ability to capture the receptor’s conformational plasticity and subtle interactions shaping ligand binding and functional selectivity.

Advances in computational modeling now provide an unprecedented view into receptor-ligand complexes. In this work, we employed AlphaFold2-based modeling pipelines (Abramson et al., 2024; Jumper et al., 2021; Varadi et al., 2022) and multistate GPCR-specific protocols (Heo & Feig, 2022) to generate high-quality structural conformations representing the inactive R, active R*, and G protein-bound R*-Gq states of *h*UT. These models were then validated against homologous GPCR data and the recently published PDB 9JFK (4 Å) structure. To probe receptor dynamics, we performed microsecond-scale molecular dynamics (MD) simulations aggregated to a total of 36 µs, which highlighted the intrinsic flexibility of transmembrane helices (TM), extracellular (ECL) and intracellular (ICL) loops, and intracellular cavity openings, all key features governing GPCR signal transduction. Finally, Site-Identification by Ligand Competitive Saturation (SILCS) (MacKerell et al., 2020) was applied to generate FragMaps that identified potential binding “hotspots” and key interaction motifs within the *h*UT binding pocket. Modeling *h*UT in complex with its endogenous ligands (R*-*h*UII and R*-URP) revealed subtle conformational shifts specific to each binding partner therefore illustrating at the molecular level their functional selectivity.

## 2. Methods

### 2.1. *h*UT modeling

The inactive (R) and active (R*) states of *h*UT, as well as the R*-G_q_ complex, were modeled as follows. The definitions of the R and R* states were based on the established GPCR activation mechanism and associated structural hallmarks described by Zhou et al. (2019). The active conformations (R* and R*-G_q_; **Figure_1 A & C**) were characterized by a 4 Å outward displacement of TM6, a 2 Å inward movement of TM5 and TM7 (**Figure_S1 A**), and an increased separation between helix 8 (H8) and the intracellular loop ICL1 (**Figure_S1 B**). A stabilizing hydrogen bond between R^3.50^ and Y^5.58^ further supported the active conformation (**Figure_S1 C**). In contrast, the inactive conformation lacked this interaction and was stabilized by a hydrophobic lock involving residues L^3.43^, I^6.40^, and V^6.41^ (**Figure_1 B** and **Figure_S1 D**). Residue numbering follows the Ballesteros-Weinstein convention (Ballesteros & Weinstein, 1995).

The amino acid sequence of *h*UT was retrieved from UniProt (accession Q9UKP6). Initial R and R* models were obtained from the GPCRdb (version 2022-08-16) (Pándy-Szekeres et al., 2018), where they were generated using AlphaFold2-Multistate (Heo & Feig, 2022). The R*-Gq heterotetramer model was constructed using AlphaFold2-Multimer via ColabFold (Abramson et al., 2024), with parameters: num_relax = 5, msa_mode = mmseqs2_uniref, model_type = multimer_v3, and num_recycles = 3. UniProt accession numbers for the G protein subunits were: Gα_q_ (P50148), Gβ (P62873), and Gγ (P59768). Canonical UniProt sequences were used in all cases.

This resulting R*-Gq complex was superimposed onto the R* model using TM1-TM4 and conserved prolines in TM5-TM7 as alignment references. Structural superposition and comparative analysis were performed using the Molecular Operating Environment (MOE) software (Chen et al., 2025). The complex was validated against homologous GPCR-G_q_ structures (PDB 7DFL and 7F6I) to confirm proper insertion of the G_q_ C-terminal α5 helix into the *h*UT intracellular cavity and its interactions with conserved residues E^3.49^, R^3.50^, A^3.53^ and R^6.32^. A GDP-Mg^2+^ complex was modeled into the Gα_q_ subunit using the GDP-bound structure PDB 7SQ2 as a template. Post-translational modifications (PTMs) were introduced using the CHARMM-GUI ‘PDB Reader and Manipulator’ module (Wu et al., 2014). These included palmitoylation of Gα_q_ Cys^9^ and Cys^10^, and geranylgeranylation of G_γ_ Cys^68^, as annotated in UniProt (T**able_S1**).

Ligand-bound active states (R*-ligand, ligands: *h*UII or URP) were generated via SILCS-Monte Carlo (SILCS-MC) sampling applied to the R* model, as described below, in subsection 2.4.

### 2.2 System construction for MD simulations

Membrane insertion and orientation of *h*UT were predicted using the PPM 3.0 server (Lomize et al., 2022), specifying the N-terminus facing the extracellular side and embedding in a mammalian plasma membrane. Protein-bilayer systems were built using the CHARMM-GUI Membrane Builder (Wu et al., 2014). Residue protonation states were assigned at pH 7 using PROPKA (https://server.poissonboltzmann.org/pdb2pqr), and annotated PTMs were included as per UniProt. These comprised N-terminal acetylation (ACE), C-terminal methylamidation (CT2), and the conserved disulfide bond between Cys^3.25^ and Cys^ECL2.50^ (**Table_S1**). To reduce system size, the N- and C-terminal regions were truncated, removing residues 1-26 and 336-389, while preserving H8 in the C-terminal segment (**Figure_S2**).

The lipid composition followed the asymmetric organization of the mammalian plasma membrane (Ingólfsson et al., 2014; Pogozheva et al., 2022; Wu et al., 2014). The upper (extracellular) leaflet included (CHARMM lipid topologies): POPC, PLPC, PAPE, POPE, SSM, NSM, N-palmitoyl-β-D-glucopyranosyl-sphingosine (GlcCer 18:1/16:0), and cholesterol in the following molar ratios of 15:21:3:3:10:10:4:35. The lower (cytoplasmic-facing) leaflet consisted of POPC, PLPC, PAPE, POPE, POPI, PAPS, SSM, NSM, cholesterol and POPA in molar ratios of 7:13:12:14:5:11:5:5:28:1. Each monomeric *h*UT systems (R, R* and R*-ligand) contained 382 lipids, while the R*-G_q_ system included 511 lipids to accommodate heterotrimer with >20 Å lateral padding. Systems were neutralized and ionized to 0.15 mM NaCl.

Final system dimensions were ≈108 x 108 x 143 Å for monomeric systems and ≈168 x 168 x 177 Å for the multimeric system. A 30 Å water layer was added above and below the bilayer, resulting in ≈ 225,000 atoms (monomers) and ≈ 307,000 atoms (multimer).

### 2.3. MD Simulation details

All MD simulations were performed with NAMD (Phillips et al., 2020) using the CHARMM36m force field (Huang et al., 2017) and TIP3P water model (Durell et al., 1994). Monomeric states (including R*-ligand models) were simulated in triplicate for 2 µs each. The R*-G_q_ system was simulated in six replicates, for a total of 36 µs. Simulations were carried out at 310.15 K under isothermal-isobaric (NPT) conditions with hydrogen mass repartitioning (Y. Gao et al., 2021) and a 4 fs time step. Temperature was maintained with Langevin dynamics (damping coefficient of 1 ps^−1^), and pressure with a Nosé-Hoover Langevin piston at 1 atm. Bonds involving hydrogens were constrained using SETTLE (Miyamoto & Kollman, 1992) for water and SHAKE (Ryckaert et al., 1977) for other molecules. Short-range cutoffs were 12 Å with a 10-12 Å switching function. Nonbonded pair lists were updated every 10 steps. Long-range electrostatics were computed with particle mesh Ewald (PME) method (Darden et al., 1993) at each integration step using a sixth-order interpolation and a grid spacing of 1 Å. System equilibration followed the CHARMM-GUI Membrane Builder protocol (Wu et al., 2014). Trajectory frames were saved every 200 ps.

### 2.4. SILCS simulations

Because both the extracellular and intracellular pockets of the R* model were accessible, this conformation was used for SILCS-based fragment mapping and docking (MacKerell et al., 2020). The R* model was embedded in a POPC:cholesterol bilayer, at a 9:1 ratio, as implemented in SILCSBio suite (MacKerell et al., 2020). The protein-membrane-fragments system was built following the standard SILCS command-line protocol for SILCS-setup (MacKerell et al., 2020) using the default fragments set, excluding halogenated fragments. Ten replicates of 100 ns GCMC/MD simulations were performed to generate FragMaps (volumetric probability maps describing the spatial distribution of pharmacophores).

### 2.5. Ligand docking with SILCS-MC

The structures of the ligands, human Urotensin II (*h*UII) and Urotensin II Related Peptide (URP) (**Figure_1 E**), were constructed from the PDB 6HVC using MOE’s Builder tool. The peptides were protonated at pH 7, and prepared for use with the CHARMM27 force field using MOE’s *Structure Preparation tool*. Ligands were energy-minimized until the RMS force gradient was below 0.01 kcal/mol/Å².

SILCS FragMap-directed docking was performed using the SilcsBio suite with the default parameters (MacKerell et al., 2020). A 25 Å radius docking grid was centered on the geometric center of *h*UT, covering all potential binding pockets but excluding extramembrane orientations. Minimization of the input structures prior to docking was enabled. Each ligand was docked in 50 replicates, each generating up to 300 poses. The 20 poses with the lowest ligand grid free energy (LGFE) values were inspected in PyMOL (The PyMOL Molecular Graphics System, Version 3.0 Schrödinger, LLC.). The best-scoring pose for each ligand was minimized in MOE within the cavity of the fixed R* model for subsequent MD system construction.

### 2.6. Trajectory analysis

#### TM helix definition

The TM helices of *h*UT were defined based on AlphaFold models as follows: TM1 (residues 45-78), TM2 (residues 88-113), TM3 (residues 121-153), TM4 (residues 166-187), TM5 (residues 206-237), TM6 (residues 256-283), TM7 (residues 291-318), and H8 (residues 322- 328).

#### Validation of Sampling Quality

Structural and sampling quality for every feature and replicate were assessed using the automated procedure implemented in PyMBAR (Chodera, 2016), which estimates a statistical inefficiency *g*, and an effective number of uncorrelated samples *N_eff_*. These diagnostics were used to confirm that our pre-selected analysis window (frames ≥5000, corresponding to the 1000 ns in the second half of each simulation) represents a conservative and statistically stable region for quantifying feature means across conditions (e.g., compare the occupancy of an H-bond in R*-URP vs R*-hUII). Replicates in which a feature was absent for the entire trajectory naturally returned no meaningful autocorrelation estimate. Such cases were treated as true biological absence, and the replicate was retained for occupancy averaging for that feature across replicates. Replicates would only have been excluded if poor sampling behavior (e.g., low effective sample size or high inefficiency), but no feature met such exclusion criteria in this study. In addition to PyMBAR-based diagnostics, overall structural stability was monitored by computing the Cα RMSD of the TM1-7 helical core and by tracking lipid-protein contact profiles (heavy atoms with a 4.5 Å contact). These metrics were evaluated to confirm that the receptor remained within a stable structural range (RMSD < 3 Å) in the analysis window used for downstream comparisons. Because GPCRs naturally explore multiple conformational microstates even in equilibrium simulations, the goal of this procedure was not to identify a strict RMSD plateau, which is not expected for these systems in simulations of this length (Latorraca et al., 2017), but rather to verify that the global fold and membrane engagement were stable during the period in which features were quantified.

#### TM-tilt measurements

Intracellular tilts of TM5, TM6, and TM7 relative to the intracellular region of TM3 were measured using the MDAnalysis (Michaud-Agrawal et al., 2011). The intracellular segment of each helix was defined as residues located in its cytoplasmic half: TM3 (residues 141-148), TM5 (residues 226-236), TM6 (residues 255-267), and TM7 (residue 313-319). For each trajectory frame, the three-dimensional center of mass (COM) of the selected Cα atoms was calculated for TM3 and for the helix of interest (TM5, TM6, or TM7). The Euclidean distance between the COM of TM3 and that of each other TM helix was then calculated, providing a time-resolved measure of intracellular displacement. Distances were stored per helix and per simulation and summarized as box-and-whisker plots generated with *matplotlib* (Hunter, 2007).

Extracellular tilts of TM5 (residues 208-215), TM6 (residues 273-279) and TM7 (residues 294-300) were measured relative to TM2 (residues 106-111) instead of TM3, as the extracellular portion of TM2 is less mobile (see **Figure_3 B**).

#### Distances separating intracellular loops

Because the loops that connect TMs have intrinsic flexibility, they can behave unpredictably, therefore, pair-wise distances were measured between intracellular loops (e.g., distance between ICL1 and ICL3) as a secondary proxy for internal cavity “closure/opening” events. MDAnalysis was used to measure the distance between the Cα atoms of residues at the extremities and middle of each loop, these were: residues 82, 84, and 86 for ICL1; 158, 160, and 162 for ICL2; 243, 246, and 249 for ICL3. For each pair of intracellular loops (ICL1-ICL2, ICL1-ICL3, ICL2-ICL3), the corresponding Cα-Cα distances were then computed for every trajectory frame, expressed in Ångströms, and stored as continuous time series. This representation captures rigid-body compaction of neighboring loops.

#### Residue flexibility

Root-mean-square fluctuations (RMSF) of Cα atoms were computed with MDAnalysis (Michaud-Agrawal et al., 2011). Each trajectory was aligned to its own average structure to remove global motions, and RMSF values were calculated as the mean deviation from time-averaged coordinates.

#### Interactions analysis

Ligand-*h*UT, intra-ligand, and intra-*h*UT interactions were identified using the *getcontacts* toolkit (https://getcontacts.github.io/) (Venkatakrishnan et al., 2019), including hydrogen bonds (sidechain-sidechain, sidechain-backbone, backbone-backbone, and water-mediated), π-stacking, T-stacking, π-cation, and salt bridges. Water-mediated interactions were recorded but omitted from figures for clarity. Only contacts persisting for at least 300 ns were reported. All data presented in this study are available in the Zenodo repository provided as supplementary material.

#### Statistical testing of contact occupancy differences

The statistical significance of differences in mean contact occupancy between R*-*h*UII and R*-URP conditions was assessed using a two-level inference framework. Within each replicate, the stationary bootstrap (Politis & Romano, 1994) with 20,000 resamples was used to estimate the sampling variance of the replicate mean while accounting for temporal autocorrelation. The mean block length was set by the optimal block length estimator of Politis and Romano as implemented in the Python *arch* package, with a minimum floor of 20 frames. Per-replicate variance estimates were then combined across the three independent replicates of each condition via inverse-variance weighting, and the difference between conditions (Δ occupancy) was tested against H₀: Δ = 0 using a Satterthwaite-approximated t-test. P-values were corrected for multiple comparisons using the Benjamini-Hochberg procedure applied within each feature type.

#### Visualization and rendering

Structural representations were generated in PyMOL v3.1.1 (the PyMOL Molecular Graphics System, Version 3.0 Schrödinger, LLC.) using color palettes selected to maximize perceptual distinctiveness under common forms of color vision deficiency. Similarly, color palettes were adequately chosen for figures generated with Matplotlib (Hunter, 2007). As PyMOL does not support shape-coding for molecular surfaces or interaction elements, color-independent, fully annotated versions of all color-encoded structural figures are provided in the Supplementary Material to ensure accessibility. To keep the figures readable, all interaction types were annotated except for hydrogen bonds (H-bond) for which the dark (blue) shade indicates sidechain to sidechain H-bonds and the light (cyan) shade indicates sidechain to backbone and backbone to backbone H-bonds, H-bonds being the most frequently detected interaction type.

## 3. Results

### 3.1. Models and Diagnostics

#### 3.1.1 Multistate models of the human Urotensin II receptor and validation

Models of *h*UT were generated in three conformational states (inactive, active and active G_q_-bound), using an approach that combines homology modeling of inactive and active GPCR templates with AlphaFold2 predictions (Jumper et al., 2021; Varadi et al., 2022), following the multi-state protocol described by Heo and Feig (Heo & Feig, 2022). The resulting models are summarized in **Figure_1** and made available in the supplementary material, while the structural features distinguishing the inactive and active states are compared in **Figure_S1**. The predicted TM topology and loop organization were consistent with canonical class A GPCR features and with previous homology models (Brancaccio et al., 2015), displaying a well-defined seven-helix TM bundle and flexible extracellular loops. Assessments of stereochemical quality (Laskowski et al., 1993) and predicted confidence scores (Studer et al., 2014) (**Table_S2 and Figure_S3**) indicated that all models satisfied standard benchmarks for high-confidence structures (Laskowski et al., 1993; Studer et al., 2014). Structural comparisons among the R and R* models revealed outward displacement of TM6 (≈ 4 Å), inward shifts of TM5 and TM7 (≈ 2 Å each), and rearrangement of H8 and the broader intracellular cavity (**Figure_S1**), consistent with activation mechanisms observed in class A GPCRs (Zhou et al., 2019).

Notably, while this manuscript was nearing completion, a cryogenic electron microscopy (cryo-EM) structure of *h*UT bound to the super-agonist P5U (PDB 9JFK; EMD-61433) was published (T. Gao et al., 2025). We retrospectively evaluated our R* model against this experimental reference. At the wwPDB-recommended contour level (0.0123), the atom-inclusion metric for the R* model is 0.91, indicating that most receptor atoms lie within regions of interpretable density (**Figure_S4 A**). A global structural superposition of R* onto 9JFK yields an RMSD of 1.291 Å, demonstrating close agreement between the modeled and experimental active conformations. Furthermore, activation microswitches PIF, CWxP, NPxxY, and DRY adopt the same rotamers and relative spatial orientations in both structures (**Figure_S4 B**). The analyses presented here rely on the multistate modeling and MD refinement pipeline available at that time. The strong agreement observed retrospectively, however, provides independent validation of the accuracy and biological relevance of the R*-based models used throughout this study.

Peptide-receptor complexes were next modeled with AlphaFold3 (Abramson et al., 2024), yielding R*-*h*UII and R*-URP complexes (**Figure_1 D**) with inter-protein TM (iPTM) scores of 0.89 and 0.95, respectively. However, the cyclic motifs of *h*UII and URP adopted identical, superimposable binding poses, indicating that AlphaFold3 did not resolve the expected differences between the two ligands. This limitation is consistent with recent reports showing that AlphaFold3 provides limited accuracy in predicting ligand-receptor complexes (He et al., 2024).

To address the uncertainty in these AlphaFold3-derived poses, we first used the SILCS method to generate three-dimensional maps (FragMaps) highlighting receptor regions energetically favorable for ligand binding (“interaction hotspots”) (section 3.2.1). These maps provided the spatial and energetic framework for subsequent SILCS-MC docking (section 3.2.2), which sampled ligand orientations within the predicted hotspots. The resulting complexes were further refined through 2 µs long-timescale MD simulations in three replicates and analyzed at the residue level (section 3.4).

#### 3.1.2. Validation of sampling quality

To validate that the analysis window of our MD simulations (the last 1 µs of each 2 µs trajectory) constitutes a well-sampled region for quantitative comparison, we conducted standard diagnostics (Chodera, 2016) that revealed statistical inefficiencies “*g*” were low and effective sample sizes “*N_eff_*” high across replicates. Importantly, no interaction feature (measured with GetContacts) showed pathological sampling behavior (e.g., low *N_eff_* or large *g*), and no replicate required exclusion when calculating average interaction occupancies (**Table_S3**). Together, these results demonstrate that structural descriptors measured over the final 1 µs provide reliable estimates of condition-dependent interaction patterns. Global structural stability metrics also show the models are behaving normally, RMSD profiles of the TM1-7 helical core remained low throughout the 2 µs simulations and never exceeded 2.2 Å relative to the starting models (**Figure_S5**). In the final 1 µs of each simulation, the average RMSD remained low : R 1.52 Å, R* 1.57 Å, R*-G_q_ 1.66 Å, R*-URP 1.52 Å, R*-*h*UII 1.65 Å. Lipid-protein contact patterns remained stable (**Figure_S6**), showing consistent profiles across replicates. These observations further confirm that the second half of each replicate simulation represents an adequate sampling regime.

### 3.2. SILCS FragMaps and Docking

#### 3.2.1. FragMaps in the active model

SILCS simulations were performed on the active-state *h*UT model embedded in a POPC/cholesterol bilayer to identify receptor regions energetically favorable for different ligand functional groups. Fragment occupancies from these simulations were converted into three-dimensional free-energy grids (“FragMaps”), which report the local solvation free energy associated with each fragment type, including water molecules.

The resulting maps revealed three dominant and spatially distinct features (**Figure_S7 A**): (i) an elongated methylamine nitrogen FragMap extending from the extracellular opening into the central cavity, indicating a pathway of favorable interactions for positively charged or hydrogen-bonding moieties; this same methylamine nitrogen FragMap also revealed a deep electrostatic hotspot near D^3.32^ in TM3, as shown in the complete set of FragMaps generated from the default fragment list (**Figure_S8**); (ii) two broad acetate carbon FragMap regions, one covering most of the extracellular cavity and loop regions above the TM bundle and another spanning the intracellular cavity enriched in lysine and arginine sidechains (these regions correspond to areas favorable for negatively charged or hydrogen-bond-accepting groups), and these acetate carbon FragMaps additionally highlighted negatively charged patches near ECL2 and at the TM7/ECL3 junction, suggesting potential coordination sites for the anionic N-terminal regions of *h*UII; and (iii) moderate-density benzene-carbon FragMap voxels distributed across the binding cavity and along membrane-exposed surfaces of the TM helices, consistent with regions accommodating hydrophobic or aromatic interactions, while the benzene carbon FragMap further delineated elongated hydrophobic and π-stacking regions proximal to aromatic receptor residues F^3.29^ in TM3, F^6.51^ and W^6.54^ in TM6, and Y^7.43^ in TM7. The complete output FragMaps directories are provided in the Supporting Information (OpenDX format).

#### 3.2.2. SILCS-MC FragMap-based docking of *h*UII and URP ligands to *h*UT

Monte Carlo FragMap-based docking (SILCS-MC) of *h*UII and URP into the precomputed FragMaps, followed by energy minimization of the top-ranked poses in the R* model, was performed as described in the Methods section. The SILCS approach quantifies ligand-receptor complementarity through the ligand grid free energy (LGFE), which reflects the chemical and spatial fit between ligand atoms and receptor environment. During SILCS-MC docking, each pose is evaluated by summing atomic LGFE contributions across the FragMaps, yielding a value (kcal.mol^-1^) that approximates the relative binding free energy within the sampled region. Lower (more negative) LGFE values correspond to stronger predicted affinities and greater physicochemical complementarity between the ligand and the receptor pocket. In SILCS-based docking, protein atoms are not explicitly considered; instead the receptor is represented by an exclusion map, which defines voxels unoccupied by fragments but occupied by protein atoms during SILCS MD simulations. Minor discrepancies between the R* model and the exclusion map occasionally produced steric clashes with receptor side-chains. Consequently, the ligand poses were subjected to energy minimization as described in Section *2.5* (*Methods*).

The LGFE scores of the ten top-ranked docking poses are reported for URP in **Table S4** and for *h*UII in **Table S5**, and the corresponding poses are illustrated in **Figure S7 B-C**. Examination of the top-ranked pose of each ligand (**Figure_2**) revealed several features consistent with previously reported structure-activity relationship (SAR) data (Brancaccio et al., 2015; Chatenet et al., 2004, 2006, 2013; Kinney et al., 2002; Sainsily et al., 2013). Notably, in both *h*UII and URP, the Lys side-chain amino group consistently occupied the deep methylamine hotspot near D^3.32^, reproducing the critical salt bridge described in mutagenesis and modeling studies (Brancaccio et al., 2015; Kinney et al., 2002; Sainsily et al., 2013). The benzene-carbon FragMaps supported the placement of each ligand’s Trp, and Tyr side chains within hydrophobic or aromatic-rich pockets, in line with experimental evidence that aromatic substitutions in URP or *h*UII strongly influence receptor binding (Chatenet et al., 2004, 2006). Minor differences emerged in the N-terminal region, where the acetate-carbon FragMap overlapped with the negatively charged Glu^1^ and Asp^4^ residues of *h*UII but not with the Ala^1^ N-terminus of URP. This observation aligns with earlier hypotheses that *h*UII’s extended acidic N-terminus (**Figure_1 E**) forms additional interactions near ECL2 (Brancaccio et al., 2015; Chatenet et al., 2013).

**Figure 2.**
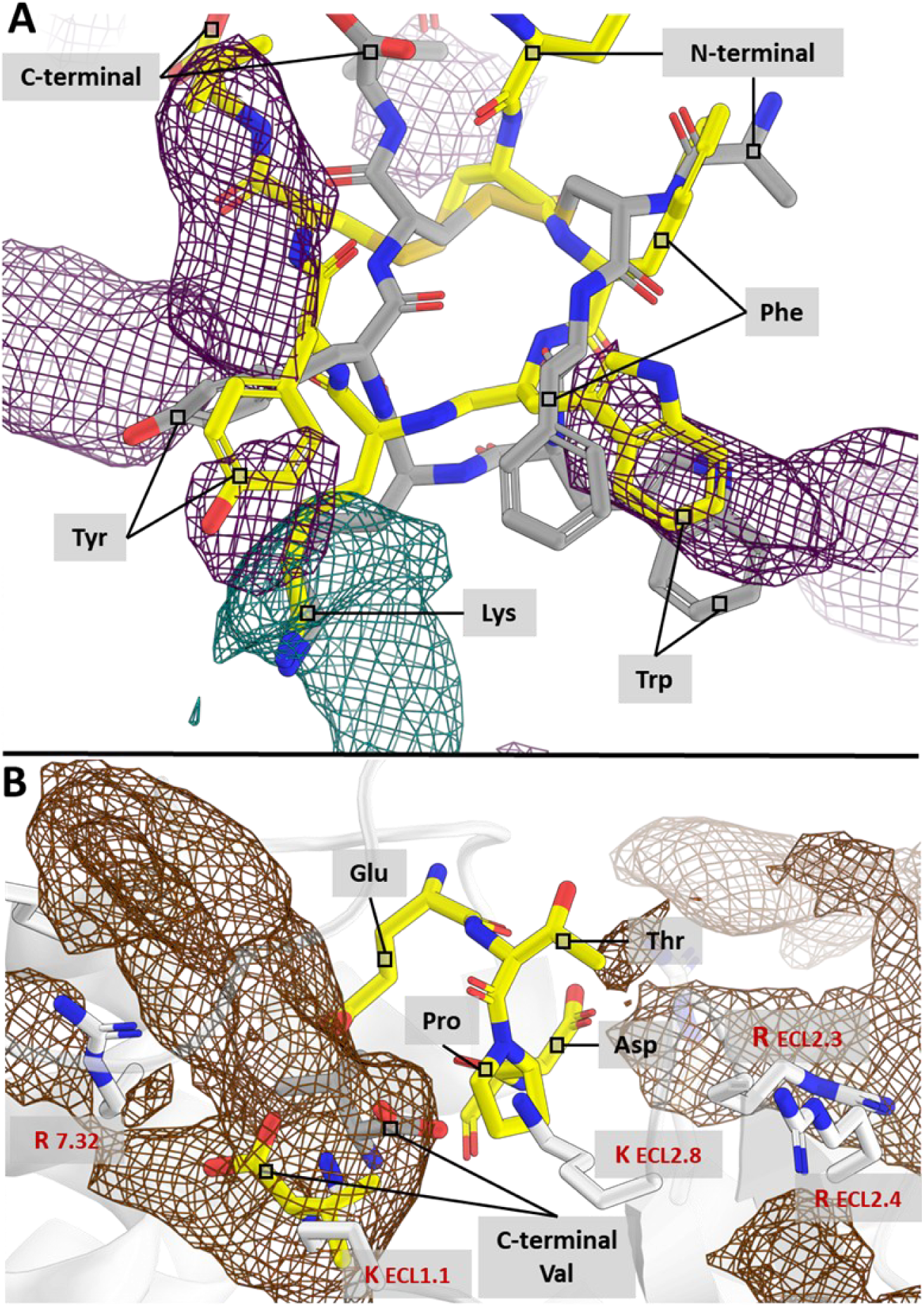
SILCS simulation of hUT in the R* state and FragMap-directed docking of hUII and URP ligands support structure-activity relationship studies reported for these peptides. (A) Top-scoring conformations from SILCS-MC docking of *h*UII (yellow carbon atoms) and URP (dark grey carbon atoms) are shown superposed with methylamine nitrogen FragMaps (blue mesh) and benzene carbon FragMaps (purple mesh). Ligand residues are labeled with their three-letter codes written in black. (B) The anionic N-terminus of *h*UII and the charged C-termini of both ligands are positioned within the extracellular pocket of *h*UT (light grey ribbon and atoms) overlaid with acetate carbon FragMaps (brown mesh) and *h*UT. Basic residues are shown as sticks (white carbon atoms) and labeled with their one-letter code and residue number in red. All hydrogens are omitted for clarity. A color-independent version of this figure is included in the Supplementary Material.

The recently published cryo-EM structure 9JFK also provides an experimental reference for the bound peptide conformation (T. Gao et al., 2025). After superposing the receptor transmembrane bundle, the backbone RMSD between the P5U ligand in 9JFK and the SILCS-MC derived poses was 0.796 Å for URP and 1.410 Å for *h*UII, indicating that both peptides adopt a binding geometry closely matching the experimentally observed active conformation (**Figure_S4 C**). In addition to this backbone similarity, the three peptides share an identical set of receptor contacts: P5U, *h*UII, and URP all have 3.5 Å contacts with residues 130, 131, 198, 280, 283, 294 and 297 in the 9JFK *h*UT structure and the R* model. Thus, both the overall peptide orientation and the residue-level contact pattern of the SILCS-MC derived complexes are consistent with the 9JFK binding mode, retrospectively supporting the use of these poses as physically realistic starting states for the subsequent MD analyses.

### 3.3. Intracellular cavity dynamics

To further characterize the *h*UT-ligand complexes, the inactive (R), active (R*), active G_q_-bound (R*-G_q_) and active ligand-bound (R*-*h*UII and R*-URP) systems were each subjected to 2-μs MD simulations, in six replicates for R*-Gq and triplicate for all other states. The final 1 µs of each trajectory were used for the analyses.

#### 3.3.1. TM-Tilt Analysis

Distances between intracellular segments of TM3 and TM5 (**Figure_S9**), TM6 (**Figure_S10**), and TM7 (**Figure_S11**) were monitored to quantify TM rearrangements and intracellular cavity opening. Figure 3A summarizes the TM5-TM3, TM6-TM3, and TM7-TM3 distance statistics. The inactive receptor exhibited a smaller TM6-TM3 distance (median ≈10 Å, Figure 3 A, model R), consistent with a “closed” conformation (Chen et al., 2025; Frei et al., 2020; Zhou et al., 2019). In contrast, R*-G_q_ displayed the largest TM6-TM3 separation (median ≈15 Å), indicative of a fully open transducer-binding cavity characteristic of active class A GPCRs (Chen et al., 2025; Frei et al., 2020). The R*-*h*UII and R*-URP systems both stabilized around 13-14 Å, with *h*UII being slightly more rigid than URP, suggesting that *h*UII constrains TM6 motion more effectively than URP relative to the unbound R* state (more on this in the section 3.4.5 below).

**Figure 3.**
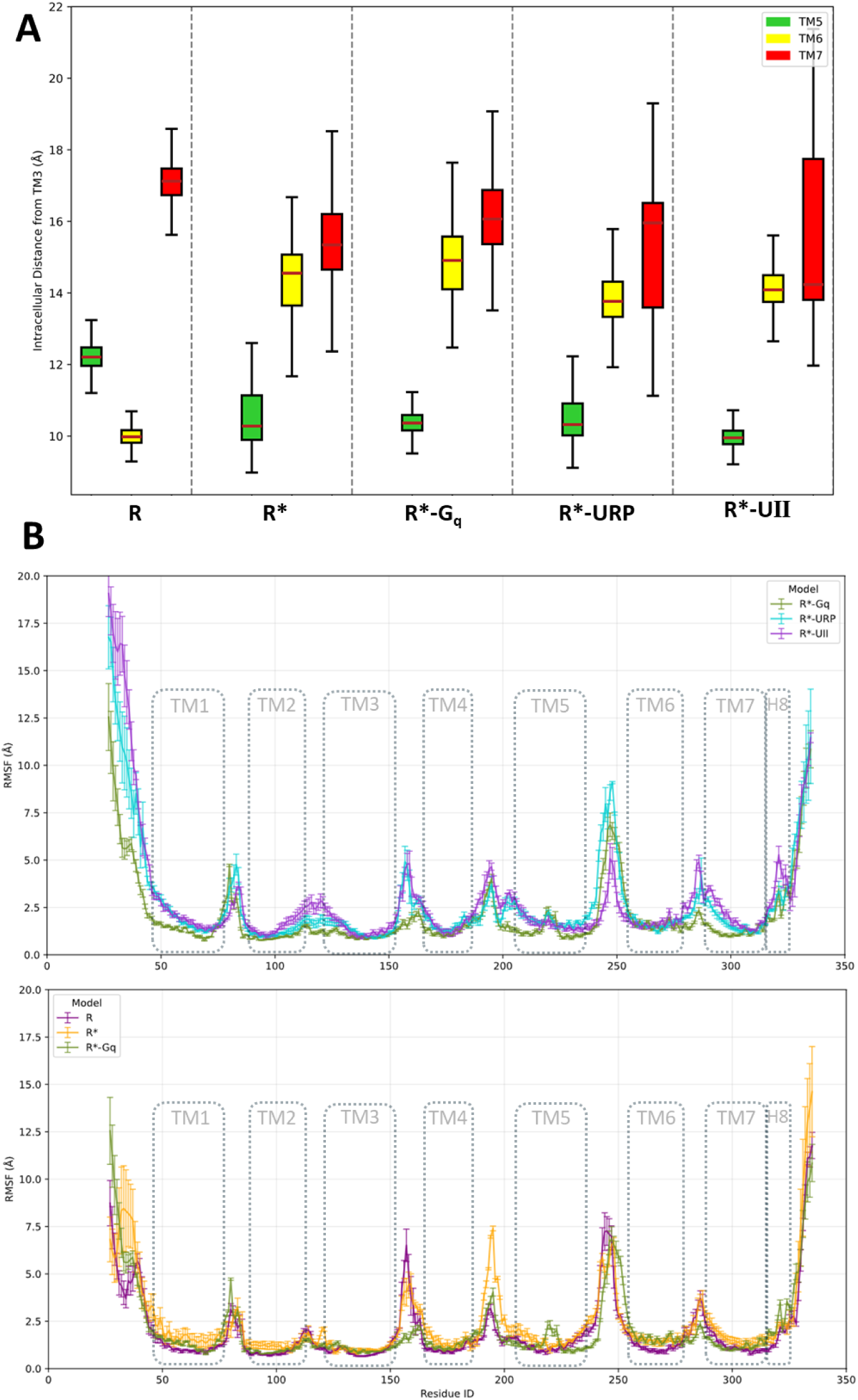
Assessment of hUT dynamics and TM-tilts that govern the accessibility of hUT’s intracellular cavity. The repartition of distances of the intracellular sections of TM5-7 to TM3 are plotted as box and whisker plots, the red line inside each box indicates the median value of each measurement, the whiskers indicate the maximum and minimum distances measured in the simulations (A). The root mean square fluctuation (RMSF) measurement across the ligand (R*-URP and R*-*h*UII) and transducer (R*-G_q_) bound simulations (B.1) and the apo simulation (B.2) plotted to compare to dynamics of the regions of interest in *h*UT. Error bars show the standard error of the mean (SEM), calculated per residue as 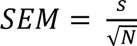 where *s* is the sample standard deviation of the RMSF values across *N* independent replicates. See paragraph *hUT TM definitions* in section *Methods* for the ranges of residues that define the TMs.

TM5-TM3 distances were generally closer in the active states (median ≈ 10-10.5 Å) than the inactive receptor (12.2 Å) (**Figure_3 A**), consistent with trends reported for other class A GPCRs (Zhou et al., 2019). R*-*h*UII again showed reduced fluctuations in TM5-TM3 distance compared with R*-URP and to the unbound R* model, while R*-G_q_ closely mirrored R*-*h*UII. TM7-TM3 distances were largest in the inactive state R (median 17.3 Å) but decreased to 14-16 Å across active states (**Figure_3 A**), a hallmark of GPCR activation (Chen et al., 2025; Frei et al., 2020; Zhou et al., 2019). Notably, R*-*h*UII exhibited the smallest median TM7-TM3 distance (14.2 Å) yet had the longest interquartile range, whereas R*-URP, R*-G_q_, and unbound R* remained near 15-16 Å.

Overall, compared with URP, the receptor bound to *h*UII shows a more rigid arrangement of TM5 and TM6 but not TM7 in the intracellular region, consistent with tighter stabilization of the active-like conformation.

#### 3.3.2. Residue RMSF Analysis of UT Flexibility

Root mean square fluctuations (RMSFs) were mapped on the receptor sequence for each simulation (**Figure_3 B**). The intracellular ends of TM5 and TM6 were notably less flexible in R*-*h*UII compared to R*-URP, reflecting *h*UII’s comparatively stronger stabilization of the intracellular cavity. TM7 exhibited the largest RMSF values in all active simulations (R*, R*-G_q_, R*-*h*UII, R*-URP), consistent with known GPCR activation processes that increase TM7 mobility. Interestingly, R*-*h*UII showed an elevated RMSF in the intracellular portion of TM7 relative to R*-URP, matching the narrower TM7-TM3 separation range noted above. These data suggest *h*UII imposes significant conformational restriction on TM5, TM6 and ICL3 (residues 238-255) but allows greater TM7-H8 positional fluctuation, whereas URP exerts less influence on TM5-TM6 (comparable to unbound R*) but results in a moderately less dynamic (lower-RMSF) TM7 intracellular tip. Additionally, both ligands seem to reduce the dynamism of *h*UT. Collectively, *h*UII’s presence seems to increase intracellular TM7 and helix 8 dynamics compared to the R*-URP, unbound R*, R*-G_q_ and R states.

### 3.4. Ligand-Receptor and Intramolecular Ligand Interactions in MD

#### 3.4.1. Global Interaction Patterns

Frames from the final 3,000 ns of each ligand-bound simulation were analyzed via the GetContacts GitHub tool to identify hydrogen bonds, salt bridges, π-π stacking, T-stacking, and cation-π interactions. Only contacts with ≥10% occupancy (i.e., present in at least 10% of the simulation) were considered (**Figure_4**). Hereafter, ligand residues will not be numbered, only *h*UT residues will be for simplicity. Consistent with prior SAR reports (Brancaccio et al., 2015; Chatenet et al., 2004, 2006), both ligands shared a central anchor: the Lys residue (**Figure_4 A & B**) forming a persistent salt bridge to D^3.32^. This Lys side chain also engaged multiple polar or aromatic residues via cation-π and hydrogen bonds, including Y^2.53^, Y^7.43^, and F^6.51^ in both complexes. The overlapping presence of these interactions corroborates the methylamine nitrogen FragMap hotspot (**Figure_2**) and earlier mutagenesis findings demonstrating the critical nature of ligand Lys for receptor binding and activation (Kinney et al., 2002; Sainsily et al., 2013).

**Figure 4.**
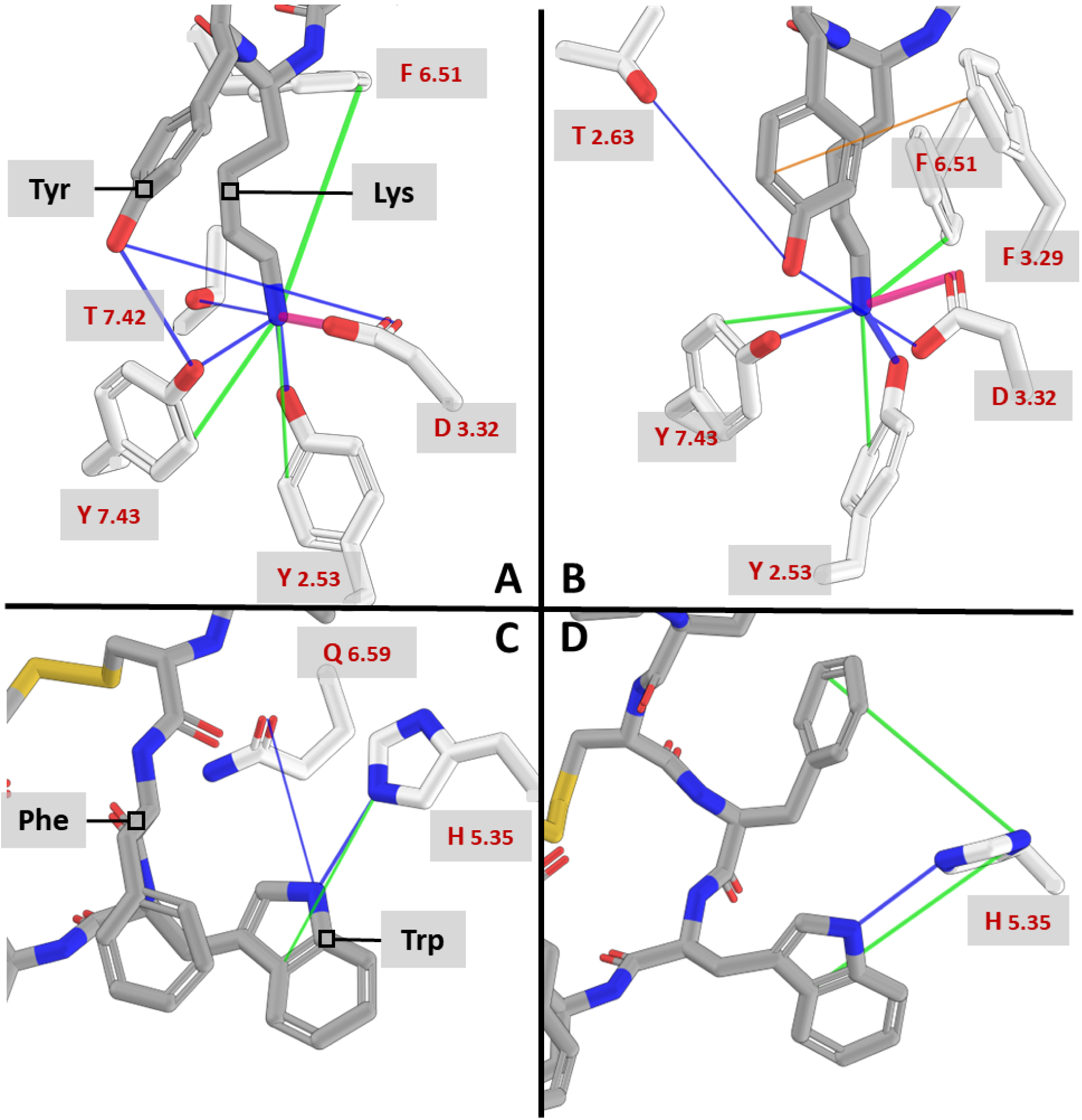
The interactions of the ligands’ cyclic core with hUT. For the R*-URP (left, A & C) and R*-*h*UII (right, B & D) simulations, interactions are represented as colored lines connecting interacting residues: sidechain hydrogen bonds (dark blue), salt bridges (magenta), cation-π (green), π-stacking (orange). Ligands are represented with dark grey carbon atoms while *h*UT residues have white carbon atoms. Interactions of the ligands Lysine and Tyrosine residues with *h*UT residues (A & B). Interactions of the ligands Tryptophan and Phenylalanine residues with *h*UT residues (C & D). All hydrogens were hidden for clarity. A color-independent version of this figure is included in the Supplementary Material.

#### 3.4.2. Differences Between *h*UII and URP Binding

Despite similar core anchoring, *h*UII and URP displayed key distinctions in aromatic side chain contacts. In URP, Tyr residue is frequently hydrogen-bonded with Y^7.43^ and D^3.32^, respectively 31% and 23% of the time in the R*-URP trajectory (**Figure_4 A**). By contrast, *h*UII’s Tyr residue showed a slightly different profile, forming hydrogen bonds with D^3.32^ (29% occurrence) and occasionally T^2.63^ (20% occurrence) rather than Y^7.43^ (7% occurrence) (**Figure_4 B**). The π-stacking interaction between Tyr and F^3.29^ occurred six-times more often in *h*UII-bound trajectories (10% occurrence) than in those bound to URP (sub-1% occurrence). Both ligands featured Trp interactions (**Figure_4 C & D**), including a hydrogen bond with H^5.35^ side-chain (with URP 35% and with *h*UII 36% occurrence) and a potential cation-π interaction (30% and 42% occurrence) with URP and *h*UII, which depends on the protonation state of H^5.35^ in the membrane microenvironment (modeled as neutral). *h*UII showed an additional potential cation-π contact between Phe and H^5.35^ occurring 47% of the trajectory time (**Figure_4 D**, see supplementary material for full interaction records), whereas this contact was absent for URP (**Figure_4 C**). These observations accord with the notion that Trp and Tyr are more critical to agonist binding affinity and receptor activation than Phe alone (Chatenet et al., 2004). In alignment with N-methylation experiments demonstrating that substitution of the backbone N-H of Phe and Tyr in URP and *h*UII similarly reduces binding affinity, but more strongly diminishes receptor activation by URP (Merlino et al., 2019), our simulations indicate that the backbone N-H of Phe and Tyr of both peptides interact with distinct regions of *h*UT. The backbone of *h*UII interacts more frequently with TM6 or TM7 residues (**Figure_5 B**) than URP (**Figure_5 A**). The same pattern was partially retained in URP (**Figure_4 A**), although with some differences in hydrogen-bond partners and frequency. These differing backbone contact patterns may contribute to the greater sensitivity of URP-based activation to backbone modification.

**Figure 5.**
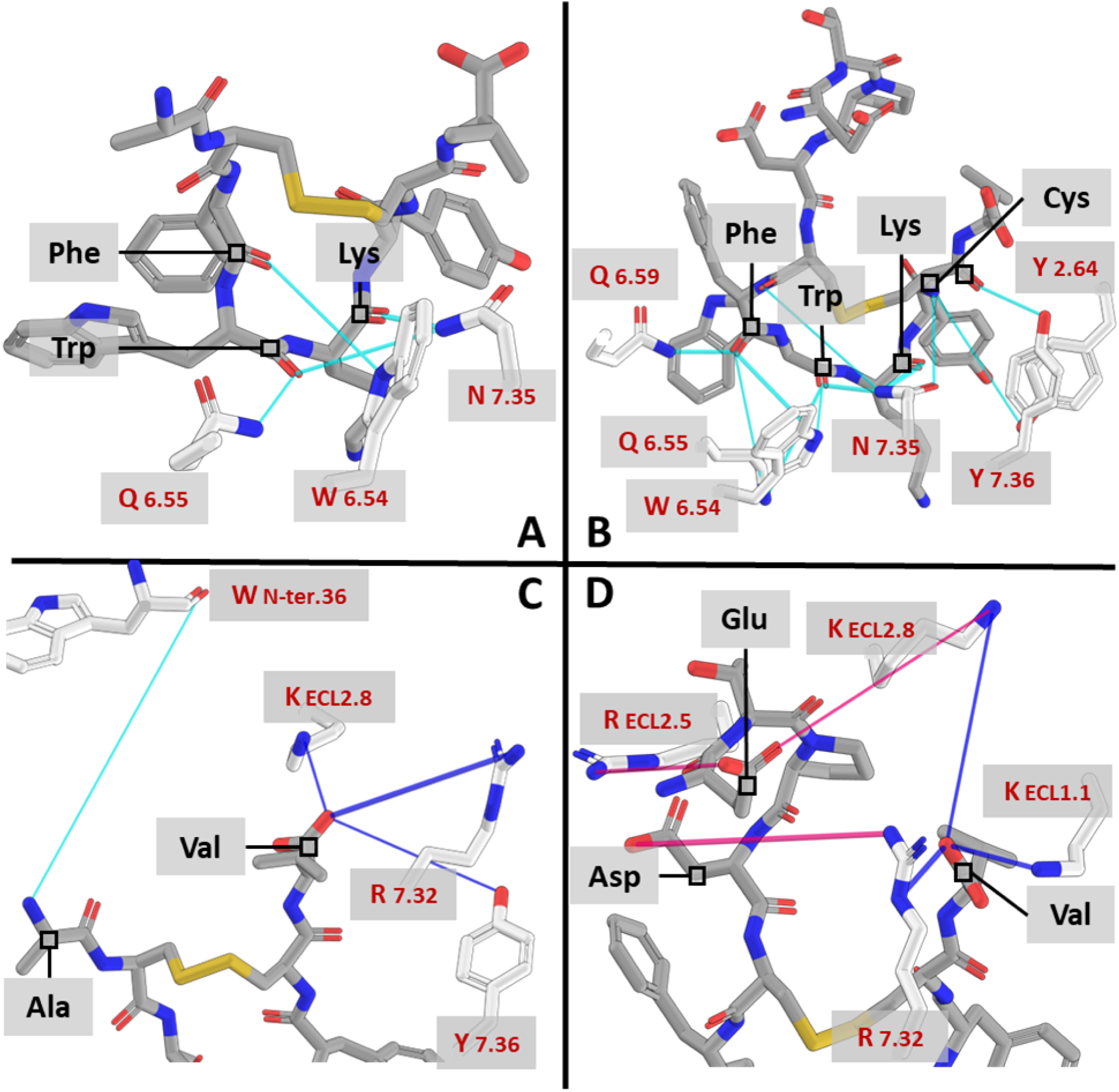
Ligand backbone and termini interactions with hUT. For the R*-URP (left, A & C) and R*-*h*UII (right, B & D) simulations, interactions are represented as colored lines connecting interacting residues: sidechain hydrogen bonds (dark blue), sidechain-backbone or backbone hydrogen bonds (light blue), salt bridges (magenta). White and dark grey carbons correspond to *h*UT and ligands, respectively. Interactions of the ligands backbone atoms with *h*UT residues (A & B). Interactions of the ligands N-terminal and C-terminal extremities with *h*UT residues (C & D). All hydrogens were hidden for clarity. A color-independent version of this figure is included in the Supplementary Material.

When analyzing the ligand-receptor interactions by calculating the difference in interaction frequencies of each receptor residue with *h*UII (from the R*-*h*UII simulation) minus its interaction frequency with URP (from the R*-URP simulation), it emerges that for those contacts shared by both complexes, *h*UII typically shows at least 10% higher frequency compared to URP. Nonetheless, URP more frequently establishes the cation-π interaction between its Lys side-chain and Y^7.43^ in *h*UT, as well as a hydrogen bond with T^7.42^, both of which are respectively less frequent and absent in the *h*UII-bound simulation.

#### 3.4.3. Divergent N-terminal and C-terminal Interactions

Major differences arose in the N-terminus, consistent with SILCS acetate Carbon FragMap predictions (**Figure_2**). *h*UII’s Glu and Asp residues were seen forming salt bridges with all extracellular loops but most notably with ECL2. In R*-*h*UII (**Figure_5 D**), salt bridges were frequently detected (> 40% of the R*-*h*UII trajectory > 1200 ns) between Glu and R^ECL2.5^ or K^ECL2.8^, along with a salt bridge between ligand Asp and R^7.32^ (upper TM7/ECL3). These interactions are consistent with earlier hypotheses that the extended anionic N-terminus of *h*UII provides additional contact points in ECLs/upper TM7 (Brancaccio et al., 2015). The neutral and small N-terminal Ala moiety of URP (**Figure_5 C**) formed a lower-frequency backbone hydrogen bond with the backbone of Trp^N-ter.36^ (in the unstructured N-terminal segment of *h*UT). This contact was evidently less stable and extensive than *h*UII’s multiple N-terminal salt bridges, aligning with the notion that URP has fewer polar interactions in the extracellular region. URP’s N-terminal is unable to establish strong interactions with extracellular loops, the only interactions we observed in the R*-URP trajectories implicating its Ala residue were weak water-bridged hydrogen bonds implicating its backbone Oxygen atom.

Both ligands carboxylate C-termini frequently interact (seen >70% of trajectory time) with K^ECL2.8^ and R^7.32^. The URP C-terminus also formed an occasional hydrogen bonding with Y^7.36^ (17% of trajectory duration), whereas *h*UII frequently involved K^ECL1.1^ (67% of trajectory duration) in bridging its carboxyl terminus (**Figure_5 C & D**).

#### 3.4.4. Intramolecular Ligand-Ligand Interactions

In both ligand simulations, intramolecular stacking was consistently observed between ligand residues Phe and Trp and between Lys and Tyr (**Figure_6 A & B**). The Phe-Tyr backbone hydrogen bond was stable in both URP and *h*UII, consistent with prior scanning studies (Merlino et al., 2019) demonstrating that N-methylation at Phe or Tyr possibly disrupts this turn-stabilizing contact which results in reduced binding and activation. This observation is compatible with the NMR-based notion that the cyclic region retains a β-turn or inverse-γ-turn conformation (Carotenuto et al., 2004; Merlino et al., 2019). These NMR-constrained models proposed that this intramolecular packing and the associated backbone hydrogen bonds stabilize the β-turn architecture essential for receptor activation (Brancaccio et al., 2015; Carotenuto et al., 2004). These differences collectively point to a finely tuned hydrogen bonding and stacking network that modulates how each ligand could maintain its core fold when bound to *h*UT. Our simulations corroborate and extend these observations within the explicit receptor environment, revealing a dynamic hydrogen-bonding and stacking network that modulates how each ligand maintains its core fold when bound to *h*UT.

**Figure 6.**
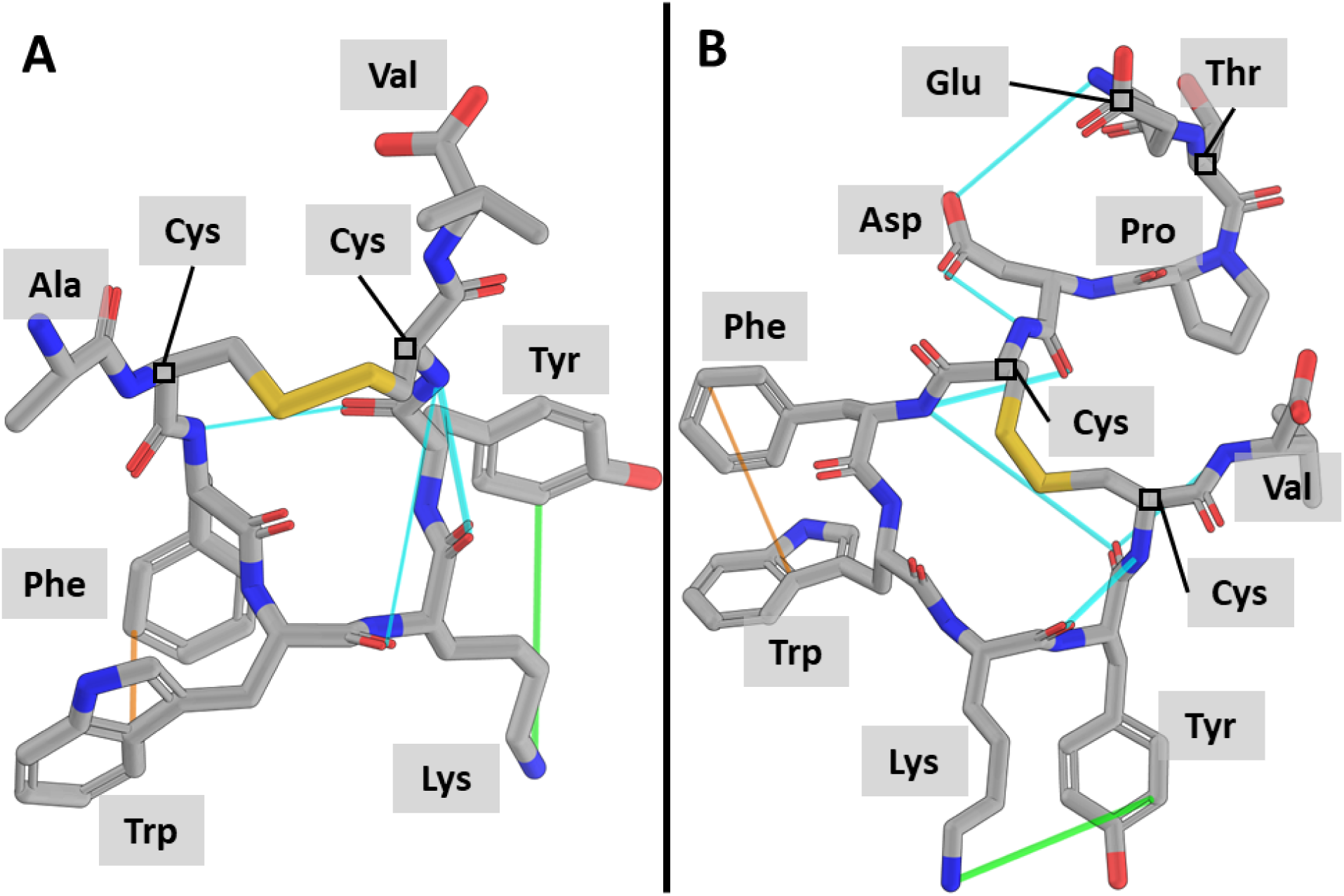
Intra-ligand interaction frequency measurements. For the R*-URP (left, A) and R*-*h*UII (right, B) simulations, intra-ligand interactions are represented as colored lines connecting interacting residues: sidechain-backbone or backbone hydrogen bonds (light blue), cation-π (green), π-stacking (orange). All hydrogens were hidden for clarity. A color-independent version of this figure is included in the Supplementary Material.

#### 3.4.5. Ligands Allosterically Induce Shifts In The Receptor’s State-Stabilizing Interactions

*h*UT is one of several class A GPCRs in which position 6.30 is not an acidic residue. Consequently, the canonical R^3.50^-E^6.30^ ionic lock that stabilizes the inactive conformation in many family members cannot form, and receptors of this subgroup are thought to populate partially active states even in the absence of agonist (Flanagan et al., 2017). Consistent with *in vitro* evidence that *h*UII and URP are both full agonists at *h*UT, the broad active-state architecture is preserved across both ligand-bound conditions, with neither ligand fundamentally rewiring the network of activation-state contacts. Rather, the most prominent ligand-dependent differences manifest as a divergence in the structural plasticity of distinct intracellular regions. Specifically, the prominent shifts are detected in the TM5-ICL3 and ICL2-TM3 junctions, and the TM7-H8 hinge, each of which adopts a different degree of contact-level order depending on which ligand occupies the orthosteric pocket.

To characterize these differences systematically, we computed the mean occupancy of every intra-protein contact detected by GetContacts in the last 1 µs of each replicate and ranked contacts by the absolute difference in mean occupancy between conditions. The top 5% contacts with the largest absolute Δ values, all mapping to the intracellular face of the receptor, are shown in **Figure_7A** and fall into three structurally coherent groups whose spatial organization is illustrated in **Figure_7B-D**.

**Figure 7.**
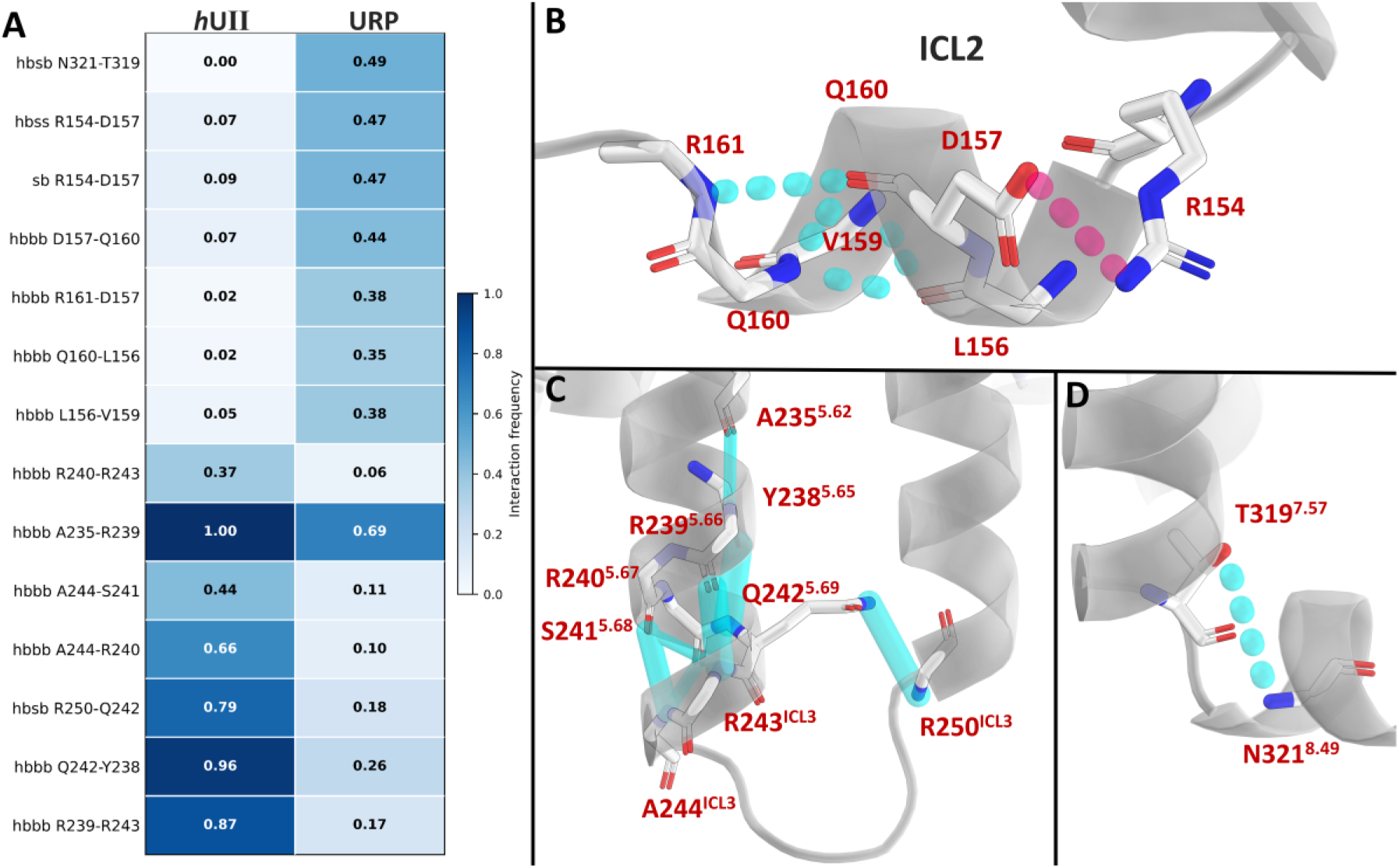
Ligand-dependent coupling of hUT cavities. (A) Heatmap of intra-protein contacts showing the largest absolute differences in mean occupancy between R*-*h*UII and R*-URP conditions (last 1 µs, triplicate simulations). Each row is one contact; left and right columns report mean occupancy in R*-*h*UII and R*-URP respectively. All contacts shown map to the intracellular face of *h*UT. Interaction type prefixes are: *hbbb*, backbone-backbone hydrogen bond; *hbsb*, sidechain-backbone hydrogen bond; *hbss*, sidechain-sidechain hydrogen bond; *sb*, salt bridge. Residue numbers follow the Ballesteros-Weinstein convention where applicable (transmembrane helices). (B) Closeup of ICL2 and the TM3-ICL2 junction (residues 154-162) in the initial R* model, with contacts enriched in the R*-URP condition shown as dotted lines or as full tubes if enriched in the R*-*h*UII condition, contact types are color encoded as cyan tubes (*hbbb*, or *hbsb*) or magenta tubes (*sb*/*hbss*). (C) Closeup of the TM5-ICL3-TM6 junction (residues 235-250) rendered as in panel B. (D) Closeup of the TM7-H8 hinge region (residues 319-321) rendered as in panel B. In all structural panels, hUT is shown as a light grey ribbon, its residues with white carbons, backbone atoms are shown without the sidechain if the latter doesn’t participate in the contact and all hydrogens are omitted for clarity. Contacts shown passed the Benjamini-Hochberg q ≤ 0.05 filter.

The seven contacts most enriched in the *h*UII-bound condition are backbone or sidechain-backbone hydrogen bonds concentrated at the intracellular end of TM5 and the first positions of ICL3 (**Figure_7 C**). The backbone hydrogen bond Q242^5.69^-Y238^5.65^ is present in 96% of R*-*h*UII frames but only 26% of R*-URP frames (Δ = +0.70). Similarly, backbone contacts in the segment flanked by residues 235-250 together define a hydrogen-bonding network that that stabilizes the TM5-ICL3 junction specifically when *h*UII is bound. Furthermore, the sidechain-backbone hydrogen bond Q242^5.69^-R250^ICL3^ (Δ = +0.61; *h*UII 0.79, URP 0.18) bridges a residue near the intracellular end of TM5 (Q242^5.69^) with a residue near the intracellular end of TM6 or start of ICL3 (R250^ICL3^), forming a cross-helix contact that couples the intracellular termini of TM5 and TM6. The persistence of this TM5-TM6 bridging contact under *h*UII reasonably provides a direct structural explanation for the tighter TM3-TM5 and TM3-TM6 intracellular distances and the lower per-residue RMSF of TM5, TM6 and ICL3 observed in R*-*h*UII simulations (**Figures_3 A-B**): by physically linking the intracellular ends of TM5 and TM6, this hydrogen bond may couple the relative tilt of both helices and reduces the conformational freedom of the loop that connects them. The backbone hydrogen bond A235^5.62^-R239^5.66^ (Δ = +0.31; *h*UII 1.00, URP 0.69), present constantly under *h*UII, reflects the greater helical regularity of the intracellular end of TM5 in this condition. Taken together, these contacts indicate that *h*UII binding is associated with a more structured, contact-rich TM5-ICL3 junction, while URP binding leaves this region comparatively plastic.

At the ICL2-TM3 boundary (**Figure_7 B**), a cluster of contacts is somewhat enriched in the URP-bound condition. The backbone hydrogen bonds in the ICL2 segment (residues 154-161) describe i+3 and i+4 backbone hydrogen bonds characteristic of an α- and a 3_10_-helical secondary structure element at the TM3-ICL2 junction. This region lies on the opposite side of the intracellular cavity from ICL3, its greater tendency to establish backbone contacts under URP is consistent with an alternative active sub-state geometry stabilized by this ligand, in which the intracellular face is organized differently from the *h*UII-bound state. However, the small occupancy values in ICL2 not exceeding 0.49 for these residues could indicate ICL2 quickly loses its helical secondary structure in the absence of the G_q_ protein contacts to stabilize the transient amphipathic ICL2 helix.

At the TM7-H8 hinge (**Figure_7 D**), the sidechain-backbone hydrogen bond N321^8.49^-T319^7.57^ is present in 49% of R*-URP frames but virtually absent in R*-hUII frames (Δ = −0.49; *h*UII 0.003, URP 0.49). This contact’s absence is a reasonable first or “piece of the puzzle” that explains the wider interquartile range of the TM3-TM7 intracellular distance (**Figure_3 A**) and why the TM7-H8 hinge is markedly more dynamic in *h*UII-bound simulations (**Figure_3 B**). Without this hinge contact, the TM7-H8 junction is free to fluctuate over a broader range of positions, producing the greater positional variance captured by both metrics.

Considered jointly, the three high-Δ contact clusters reveal that *h*UII and URP impose complementary but spatially separated patterns of structural order on the intracellular face of *h*UT. *h*UII stabilizes the TM5-ICL3 junction and the cross-helix TM5-TM6 bridge while leaving the TM7-H8 hinge unconstrained, whereas URP stabilizes the ICL2-TM3 boundary and the TM7-H8 hinge while leaving the TM5-ICL3 junction structurally plastic. The mechanistic link between the orthosteric binding differences documented in chapter 3 and these intracellular rearrangements remains allosteric in nature and cannot be resolved from the present simulations alone; however, the spatial coherence and magnitude of the observed contact differences (Δ up to 0.71 across three independent replicates) establish that the two ligands likely reorganize the intracellular face of *h*UT in distinct ways despite their near-identical binding poses at the orthosteric site.

## 4. Discussion

This study presents a comprehensive integrative modeling strategy to resolve the structural and dynamic nuances of the urotensin II receptor interactions with its two endogenous ligands, *h*UII and URP. By bridging prepared AlphaFold-derived multistate models, SILCS FragMap based docking, and microsecond-scale molecular dynamics simulations, we provide new insights into the structural determinants of ligand-specific activation within the urotensinergic system.

The SILCS FragMaps identified physico-chemically distinct sub-pockets within the transmembrane binding cavity, many of which recapitulate known structure-activity relationship data for both *h*UII and URP (Chatenet et al., 2004, 2006, 2013; Kinney et al., 2002; Merlino et al., 2019; Sainsily et al., 2013). The high frequency occupancy of methylamine nitrogen and benzene carbon FragMaps in regions encompassing D^3.32^, F^6.51^, and Y^7.43^ validates prior mutagenesis and pharmacophore studies and supports the robustness of the model in identifying pharmacologically relevant interaction hotspots (Brancaccio et al., 2015; T. Gao et al., 2025; Kinney et al., 2002; Sainsily et al., 2013). These data, together with the fragment-directed SILCS-MC simulations, reinforce the central role of the Lys-D^3.32^ salt bridge as a conserved anchoring mechanism while uncovering subtle, ligand-specific variations in polar and aromatic side-chain engagement, especially involving ECL2 and TM6-TM7 residues. The top-ranked poses (lowest LGFE score) for *h*UII and URP (**Tables S4 and Table S5**) were also consistent with the docking solutions proposed by Brancaccio *et al*. (2015) and the recent cryo-electron microscopy structure of *h*UT bound to its ligand P5U (PDB: 9JFK, published during time of writing) (T. Gao et al., 2025), confirming that the Lys-D^3.32^ salt bridge is pivotal for ligand-receptor complexes.

Our long-timescale simulations further suggest that these shared anchoring mechanisms can produce divergent dynamic consequences. While both ligands stabilized the receptor in an active-like conformation, *h*UII imposed more pronounced constraints on the intracellular regions of TM5 and TM6, effectively reducing the mobility of the transducer-binding cavity potentially influencing subsequent ligand-induced signal transducer recruitment. URP, although similarly bound to D^3.32^, permitted greater flexibility in TM6 and TM7, suggesting a more permissive and potentially kinetically distinct active conformation. The differences observed in the degree of structural order at the TM5-ICL3 junction, the ICL2-TM3 boundary, and the TM7-H8 hinge (**Figure_7**) further support the hypothesis that ligand structural features outside the core pharmacophore, such as the acidic N-terminal extension in *h*UII, may fine-tune receptor activation by engaging distinct microdomains at the extracellular face.

From a drug discovery perspective, these findings highlight several opportunities. The fact that SILCS FragMaps not only mirrored but extended established SAR patterns suggests that fragment-based hotspot mapping can serve as a powerful complement to conventional pocket identification in GPCR systems, especially for pocket selective ligands. Furthermore, the observation that small differences in ligand structure and interaction translate into distinct stabilization patterns for transmembrane helices suggests a mechanistic basis for biased signaling, providing a molecular rationale for selective pathway engagement. These insights can inform the design of new agonists that exploit peripheral interactions, particularly in the extracellular vestibule and helix interfaces, to steer receptor activity towards β-arrestin-biased outcomes by stabilizing a G protein-incompetent state, a pathway increasingly associated with cardioprotective effects on *h*UT. More concretely, our simulations delineate two recurrent interaction fingerprints that can be used as design templates: an “*h*UII-like” class, characterized by an acidic N-terminal motif engaging ECL2 and R^7.32^ to draw TM5-TM6 inward, dense aromatic and cation-π contacts centered on F^3.29^ and H^5.35^; and a “URP-like” class, in which a compact, neutral N-terminus interacts only weakly with the extracellular loops, ligand Lys and Tyr residues redistribute their contacts toward Y^7.43^/T^7.42^, and TM5-TM6 remain more dynamic. Synthetic agonists can be steered along this continuum by introducing or removing acidic groups in pocket delineated by the extracellular loops, repositioning aromatic residues to favor F^3.29^/H^5.35^ versus T^7.42^/Y^7.43^ contacts, and modulating backbone hydrogen-bonding capacity within the cyclic core, thereby providing a physically grounded handle on receptor-state selection while preserving the shared Lys-D^3.32^ anchoring pharmacophore. In this sense, the *h*UII-like and URP-like templates illustrate how modest changes in peripheral and backbone interactions can rebalance the relative stabilization of partially active versus fully active conformations, and thus modulate the timing, amplitude, and pathway preference of downstream signaling rather than enforcing a strictly “β-arrestin-only” profile. We therefore propose that exploiting these interaction fingerprints offers a route to tune functional selectivity at *h*UT, for example by enriching signaling signatures associated with coordinated G protein and β-arrestin engagement and cardioprotection, without departing from the endogenous ligand scaffold.

Compared with earlier studies that relied on NMR-derived ligand conformations embedded in detergent micelles or SDS environments and subsequently docked into rigid or partially flexible receptor models, our combined SILCS and MD approach provides a more physically grounded representation of the binding process. The fragment-based SILCS sampling captures the complete three-dimensional free-energy landscape of the *h*UT pocket in explicit solvent and membrane, thereby integrating local desolvation effects that strongly influence ligand binding. Subsequent long-timescale molecular dynamics simulations propagate these energetically preferred poses within a dynamic receptor framework, allowing the simultaneous quantification of both ligand-receptor and receptor-intrinsic state-stabilizing interactions. This workflow therefore bridges the gap between static structures and true dynamical behavior: it recovers the experimental structure-activity relationships established by earlier NMR-based models and matches the structural accuracy of the recently published *h*UT cryo-EM structure but extends them by revealing how ligand-specific hydrogen-bonding, stacking, and electrostatic networks shape GPCR intracellular contact networks. In doing so, our approach unifies sampling of receptor conformational dynamics with quantitative interaction profiling, offering a more predictive and mechanistically faithful route to dissect functional selectivity within the urotensinergic system.

In conclusion, the molecular portrait developed here resolves how the two closely related peptide ligands can differentially influence conformational dynamics and stabilization patterns in *h*UT. By demonstrating that ligand-specific dynamics emerge not only from core anchoring interactions but also from peripheral loop and helix contacts, this study provides a conceptual and methodological framework for advancing *h*UT-targeted drug discovery and uses methods that can deepen our understanding of functional selectivity in class A GPCRs.

## 5. Conclusion

The data presented here establish that the AlphaFold Multistate-derived *h*UT models behave in a manner consistent with established GPCR activation motifs, previously described *h*UT-ligand structure-activity relationships and the recently published cryo-EM structure of *h*UT. SILCS FragMaps corroborate the central importance of ligand Lys-D^3.32^ salt bridging and aromatic side chain burial, and the SILCS-MC docking simulations reproduce prior homology-based poses (Brancaccio et al., 2015), a 4 Å cryo-EM structure (T. Gao et al., 2025), as well as SAR findings (Chatenet et al., 2004, 2006, 2013; Kinney et al., 2002; Merlino et al., 2019; Sainsily et al., 2013). Long-timescale MD simulations highlighted that *h*UII exerts a stronger constraint on TM5 and TM6, stabilizing an active-like receptor conformation, whereas URP leaves TM5 and TM6 more flexible, stabilizing an alternative active-like conformation. Distinctions also arise at the receptor’s extracellular loops, where the extended acidic N-terminal region of *h*UII engages ECL2 more extensively than the simpler Ala-based N-terminus of URP. These differences collectively suggest that *h*UII and URP share a conserved anchoring mechanism but diverge in how they stabilize the *h*UT receptor’s transmembrane and extracellular regions, potentially influencing differential kinetics and functional selectivity in different receptor microenvironments.

Altogether, these integrative methods enabled us to gain insights into how *h*UII and URP engage distinct receptor microdomains and stabilize alternative active conformations of *h*UT thus clarifying structure-function relationships that drive intracellular rearrangements potentially favoring different downstream signal transducers. By capturing the dynamic range of *h*UT conformations and mapping ligand interactions, our study contributes with a valuable tool for rational atomic-scale drug design.

## Glossary

*h*UII (human Urotensin II peptide), URP (Urotensin II related peptide), *h*UT (human Urotensin receptor), GPCR (G protein coupled receptor), SAR (structure activity relationship), TM (transmembrane helix), H8 (helix 8), ECL (extracellular loop), ICL (intracellular loop), MD (molecular dynamics), PTM (post-transcriptional modification), FragMaps (3D fragment-occupancy probability density map), SILCS (site identificiation by ligand competitive saturation), MC (Monte-Carlo sampling of ligand conformational space), LGFE (ligand grid free energy).

## Associated Content

The Supporting Information will be available on Zenodo 10.5281/zenodo.15713368 :

- Apo R*, R and R*-Gq AlphaFold models;
- Top 10 receptor-ligand poses;
- Simulated MD systems PDB and PSF files;
- MD simulation DCD files saved at 10 ns per frame;
- Interactions full record.

## Acknowledgments

For providing access to their computational resources the author thanks the professors David Chatenet from the Institut National de la Recherche Scientifique and Patrick Lagüe from the university of Laval, Quebec, Canada. This research was enabled in part by computational resources provided by Calcul Québec (calculquebec.ca) and the Digital Research Alliance of Canada (alliancecan.ca).

## Author contributions

A.G.T. designed the study, performed the experiments, analyzed the data, and wrote the original manuscript.

