## Supplemental Figures & Tables for "Integrative AlphaFold Modeling, Fragment Mapping, and Microsecond Molecular Dynamics Reveal Ligand-Specific Structural Plasticity at the Human Urotensin II Receptor"

Supplementary Information

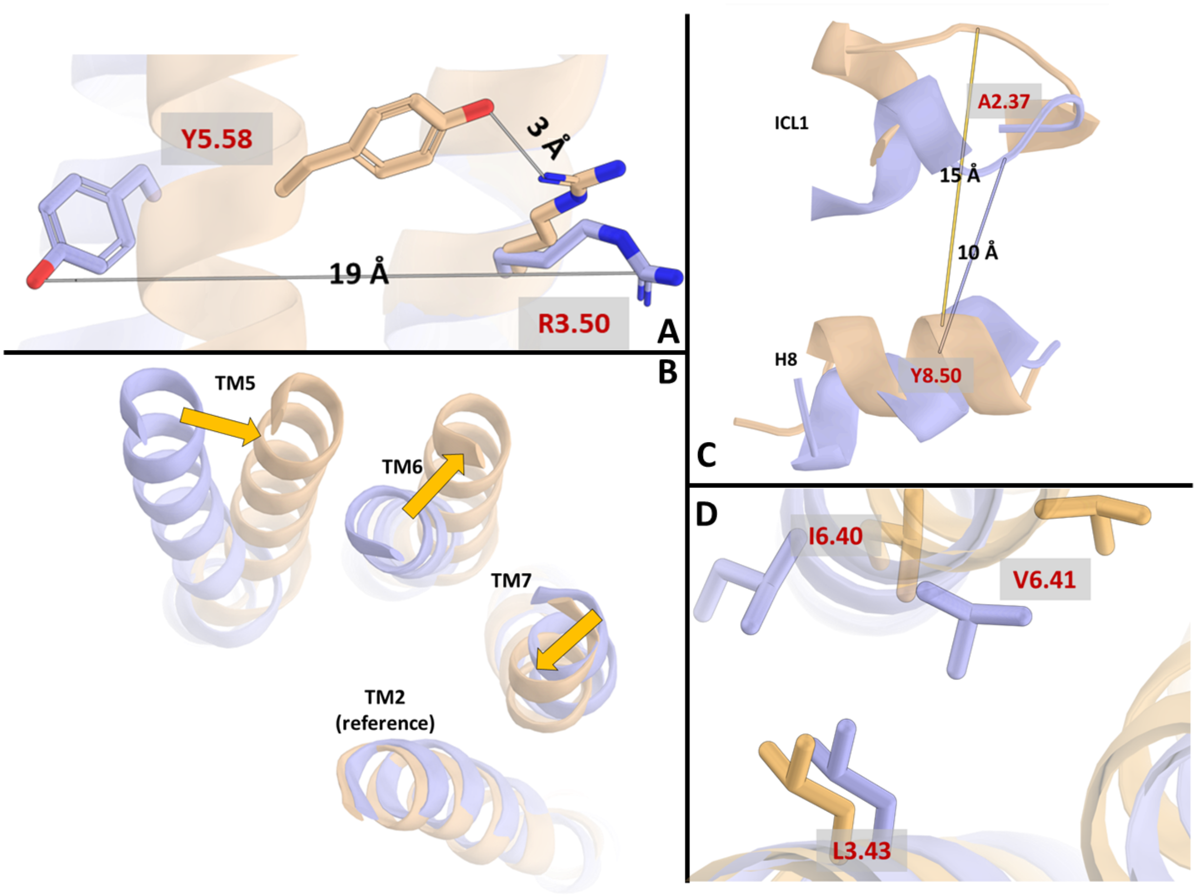


**Figure_S1**. Comparison between the modeled inactive (R, light blue) and active (R*, light orange) states of *h*UT. The models exhibit the characteristic hallmarks of GPCR activation, including the slight inward tilt of TM5 on the intracellular side, which enables the formation of the stabilizing $R^{3.50}$-$Y^{5.58}$ interaction (panels A and B). Activation is further accompanied by the outward displacement of TM6 and inward tilt of TM7 (panel B), which together shift H8 away from the intracellular loops. This motion, measured as the distance between the Cα atoms of $A^{2.37}$ and $Y^{8.50}$, reduces H8 contacts to ICL1 (panel C, distances in blue and orange correspond to the R and R* states, respectively) and disrupts the hydrophobic lock formed by the residues $L^{3.43}$, $I^{6.40}$, $V^{6.41}$(panel D).

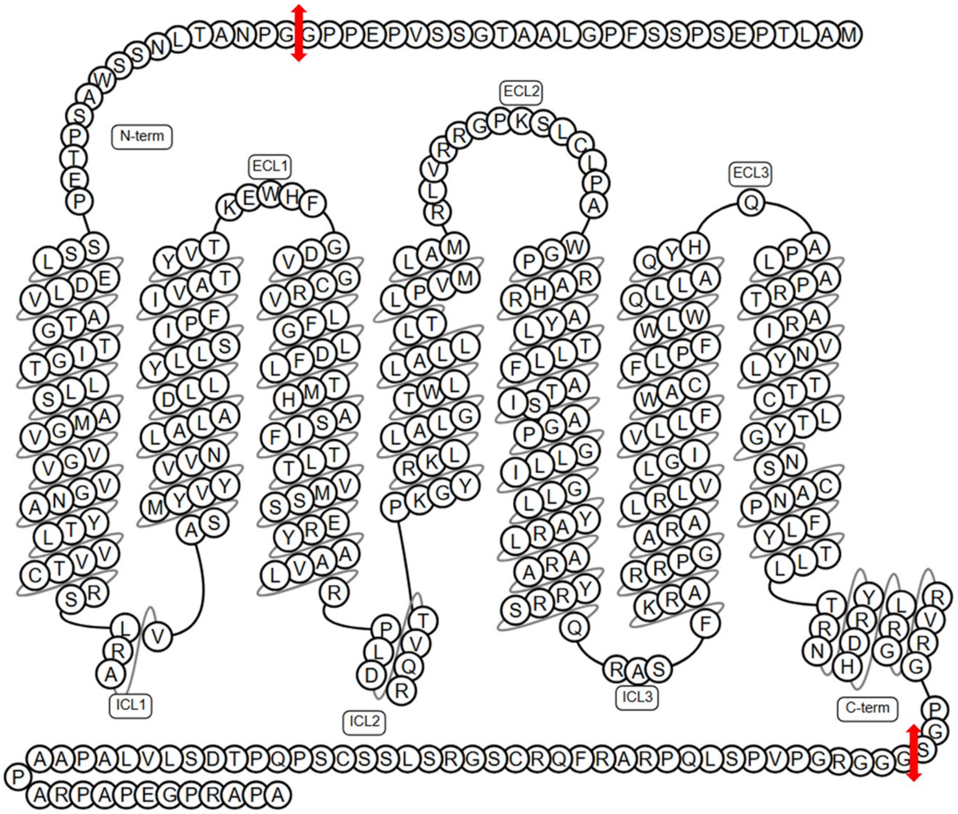


**Figure_S2**. Truncations of *h*UT terminal segments removing residues 1-26 and 336-389, indicated by two red arrows, were made to minimize the required MD system size. H8 in the C-terminal was preserved.

**Table S1**. List of all post-translational modifications (PTMs) and structural preparation applied to the *h*UT and G-protein models. PTMs marked as 'NA' indicate that no modification was applied to the corresponding protein. No phosphorylation sites were included.

| | **UT** | **Gαq** | **Gβ** | **Gγ** |
|---|---|---|---|---|
| **Terminal truncation (Residues)** | 1-26 and 336-389 | NA | 1 (methionine) | 1 (methionine) |
| **Disulfide bond** | C123-C199 | NA | NA | NA |
| **Acylation** | NA | C9 and C10 (S-palmitoylation) | NA | S-geranylgeranylation on C68 |
| **Protonation (PROPKA) at pH** | 7 | 7 | 7 | 7 |

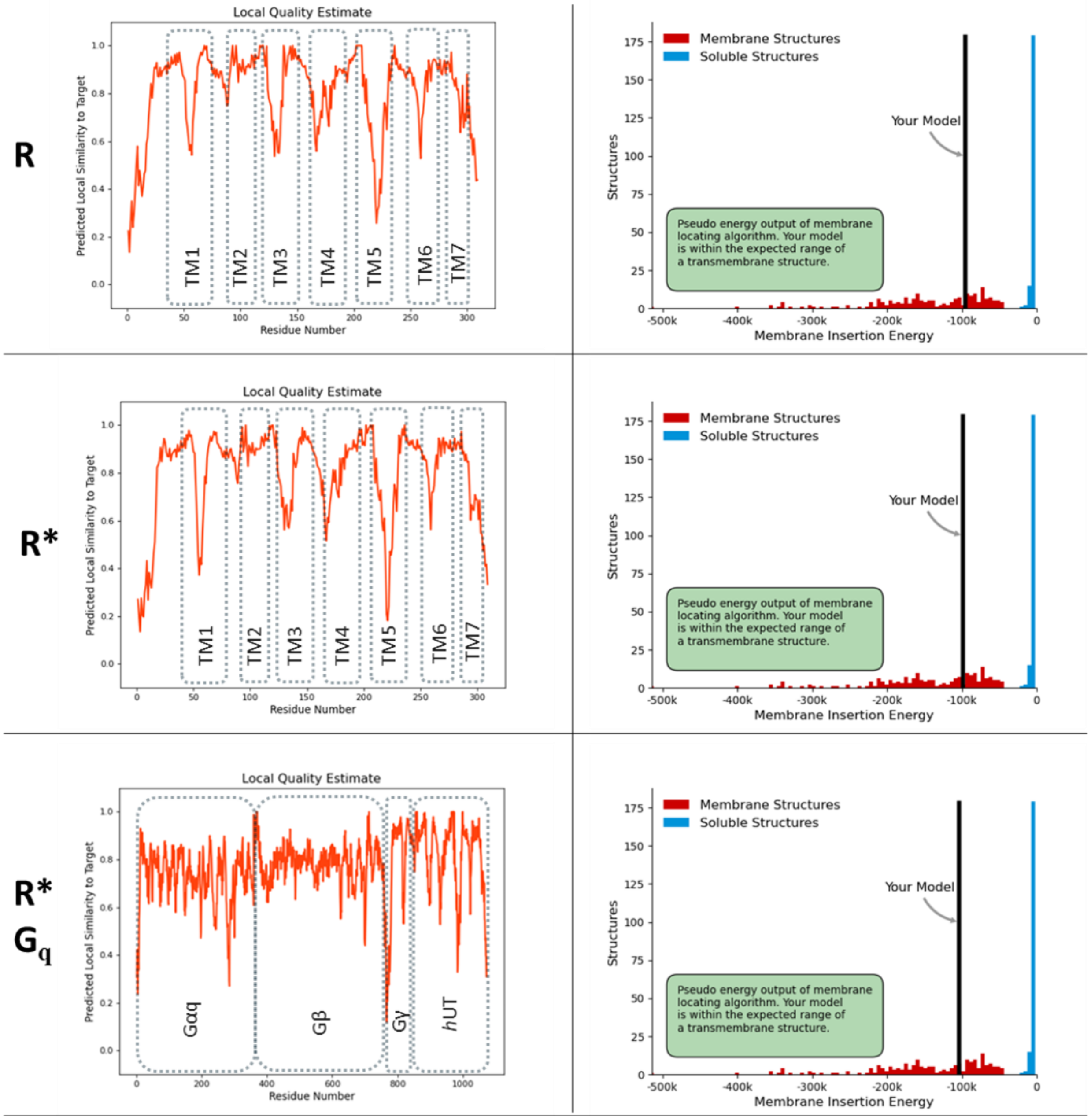


**Figure_S3**. Quality assessment of the *h*UT models on SwissModel-QMEANBrane webservice (https://swissmodel.expasy.org/qmean/) [add ref9]

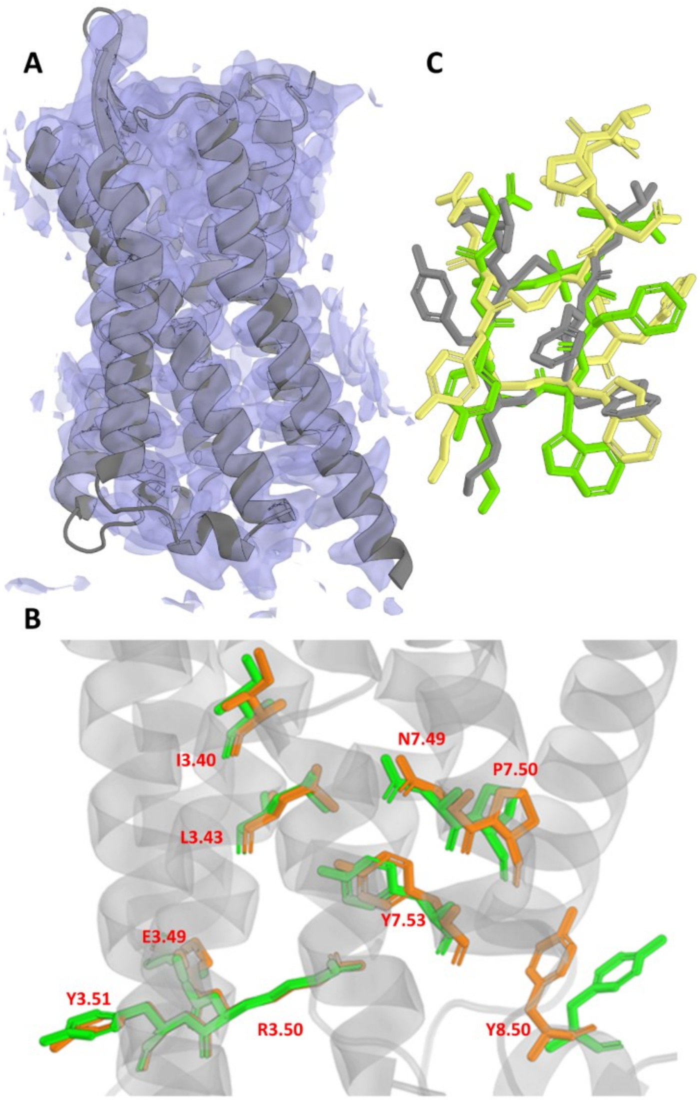


**Figure_S4**. Validation of the R* model and SILCS-MC peptide poses using the *h*UT-P5U cryo-EM structure (9JFK). The active-state receptor model (R*) displayed in the experimental cryo-EM electron density map (EMD-61433) at the recommended contour level 0.0123 (panel A). The model fits the density across the transmembrane bundle, consistent with an atom-inclusion score of 0.91. Superposition of the R* model with the experimentally resolved *h*UT receptor from 9JFK, key activation microswitch residues from the PIF, NPxxY, and DRY motifs and conserved residue 8.50 are shown as sticks (orange, R*; green, 9JFK), highlighting the close agreement in side-chain rotamers and spatial orientations between the modeled and experimentally observed active conformations (panel B). Overlay of the three peptide ligands after aligning the R* receptor to the 9JFK structure. The SILCS-MC predicted poses of the URP (grey) and *h*UII (yellow) peptides adopt backbone conformations closely matching the experimental P5U ligand (green), with backbone RMSDs of 0.796 Å (URP) and 1.410 Å (*h*UII). All three peptides share the same receptor-facing orientation supporting the accuracy and biological relevance of the SILCS-derived binding poses used in the subsequent simulations R*-URP and R*-*h*UII.

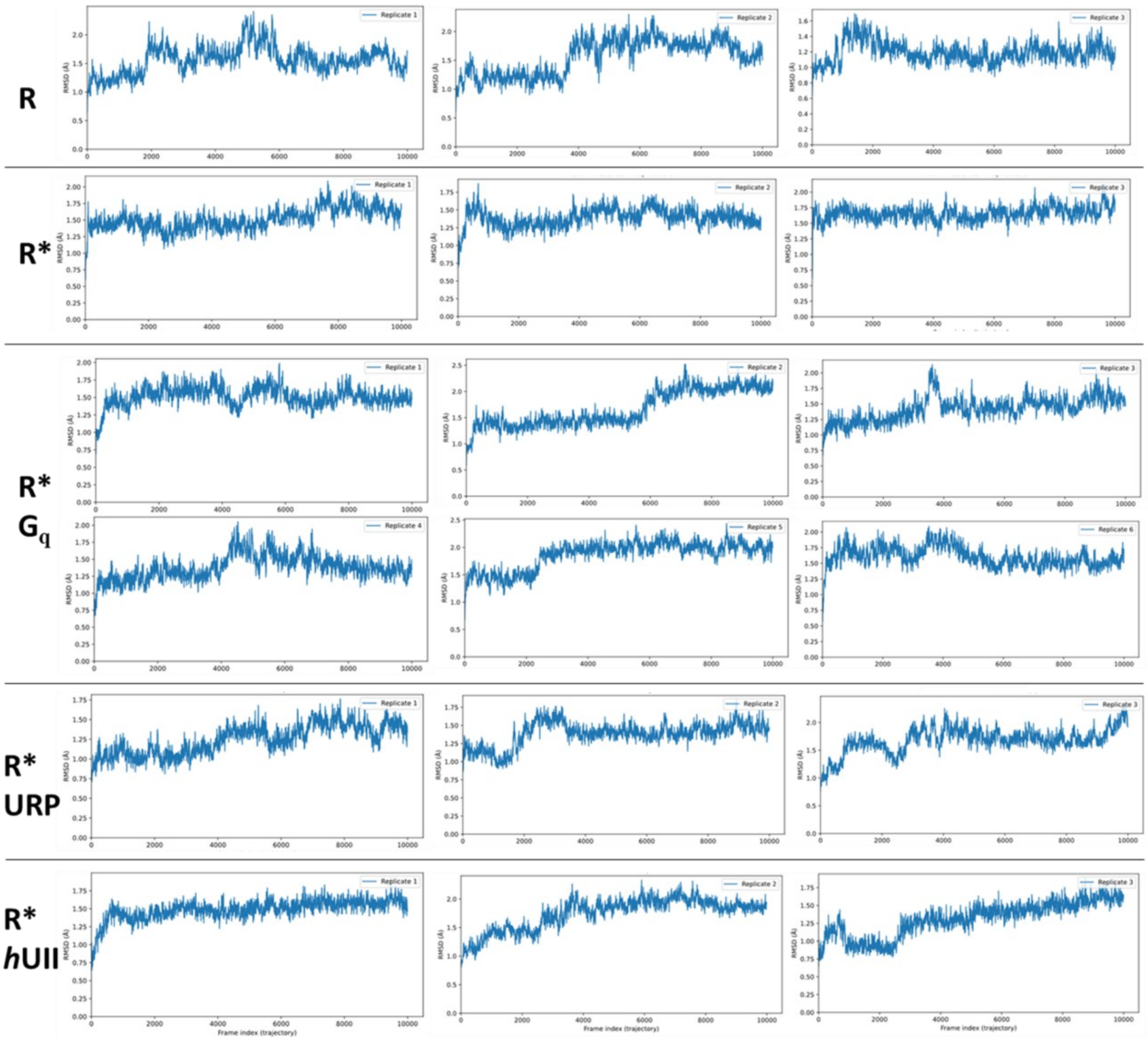


**Figure_S4**. RMSD time series of each replicate simulation, frames were saved at a 200 ps interval, thus 10,000 frames correspond to the full 2000 ns trajectories. RMSD was computed on alpha-carbon atoms of *h*UT TM1-7 residues. See *Methods* section for *h*UT TM definition. The average RMSD values in the second half of each replicate (last 5000 frames corresponding to 1000 ns) are as follows: for the state **R** 1.52 Å, **R*** 1.57 Å, **R*-G$_q$** 1.66 Å, **R*-URP** 1.52 Å, **R*-*h*UII** 1.65 Å, thus considering the systems stable while they explore the multiple conformational basins known for GPCRs.

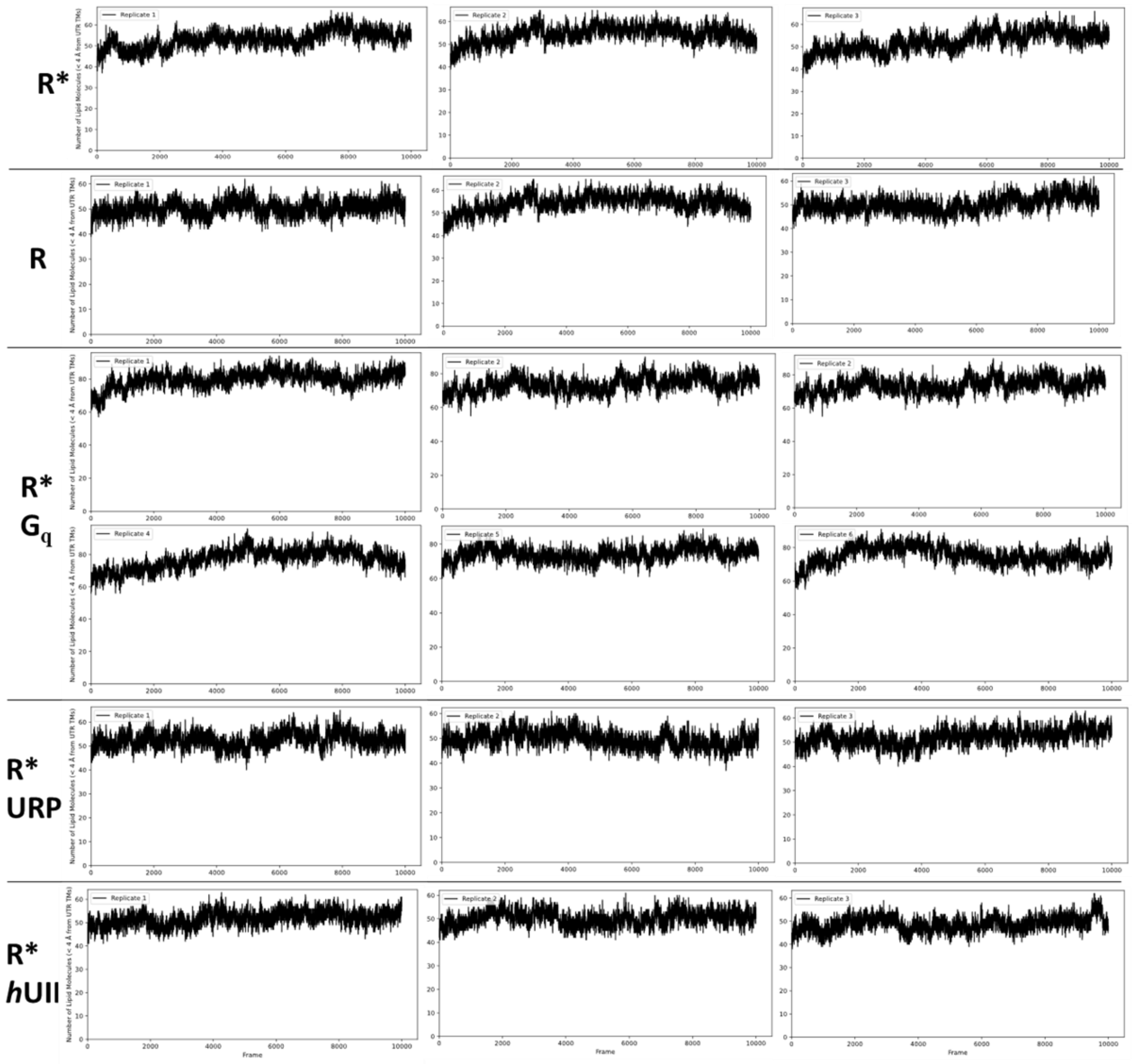


**Figure_S5**. Lipid molecule-*h*UT contact time series of each replicate simulation, frames were saved at a 200 ps interval, thus 10,000 frames correspond to the full 2000 ns trajectories. Contacts were recorded as a lipid molecule having at least one heavy atom within 4 Å of any *h*UT TM residue heavy atom. The Y axis represents the number of lipid molecules that have a contact with *h*UT TM residues only excluding highly flexible loops. See *Methods* section for *h*UT TM definition.

**Table_S2**: Quality assessment of *h*UT models was performed using PROCHECK using the SAVES v6.1 server (https://saves.mbi.ucla.edu/) [ref add8]. Numbers in parentheses indicate the count of residues classified as Ramachandran outliers, all located in unstructured loops near proline residues.

| *h*UT model | Ramachandran favored (%) | Rotamers favored (%) |
|---|---|---|
| R | 100 | 100 |
| R* | 99.6 (1) | 100 |
| R*-$G_q$ | 99.3 (7) | 100 |

**Table_S3**: Sampling diagnostics for GetContacts interaction features. For the simulations of R*-URP and R*-*h*UII, the table reports: total number of analyzed interactions (the higher number for *h*UII is due to its longer N-terminal sequence); the percentage of interactions with $N_{eff} > 500$ samples (sufficient sampling); the median statistical inefficiency *g*; and the fractions of features with mean occupancy ≥ 0.1, < 0.1, or absent. These metrics summarize MD sampling quality and interaction prevalence in the analysis window. All conditions showed uniformly high Neff and low g, indicating adequate decorrelation and reliable interaction occupancy estimates.

| Condition | Number of interactions | $N_{eff} > 500$ | Median *g* | Occupancy ≥ 0.1 | Occupancy < 0.1 | Absent |
|---|---|---|---|---|---|---|
| R*-URP | 60 | 100% | 1.15 | 49.4% | 35% | 15.6% |
| R*-*h*UII | 83 | 100% | 1.38 | 46.6% | 32.5% | 20.9% |

**Table_S4**: Scores of the ten top-ranked URP poses obtained from SILCS-MC docking. The total energy (TotE) and the ligand grid free energy (LGFE) are reported in kcal mol$^{-1}$, the ligand efficiency (LE) in kcal mol$^{-1}$ per heavy atom.

| Pose ranking | TotE | LGFE | LE |
|---|---|---|---|
| 1 | 36.516 | -10.055 | -0.142 |
| 2 | 36.467 | -9.572 | -0.135 |
| 3 | 33.945 | -9.569 | -0.135 |
| 4 | 34.825 | -9.38 | -0.132 |
| 5 | 34.313 | -9.326 | -0.131 |

| | | | |
|---|---|---|---|
| 6 | 34.073 | -9.26 | -0.13 |
| 7 | 35.343 | -9.147 | -0.129 |
| 8 | 34.144 | -9.126 | -0.129 |
| 9 | 33.988 | -9.045 | -0.127 |
| 10 | 33.943 | -8.996 | -0.127 |

**Table_S5**: Scores of the ten top-ranked *h*UII poses obtained from SILCS-MC docking. Units are identical to those in Table S4.

| Pose | TotE | LGFE | LE |
|---|---|---|---|
| 1 | 61.273 | -11.403 | -0.118 |
| 2 | 59.92 | -9.731 | -0.1 |
| 3 | 60.329 | -9.151 | -0.094 |
| 4 | 56.651 | -9.097 | -0.094 |
| 5 | 53.85 | -9.021 | -0.093 |
| 6 | 61.229 | -8.956 | -0.092 |
| 7 | 56.572 | -8.892 | -0.092 |
| 8 | 61.388 | -8.797 | -0.091 |
| 9 | 58.141 | -8.053 | -0.083 |
| 10 | 58.966 | -8.033 | -0.083 |

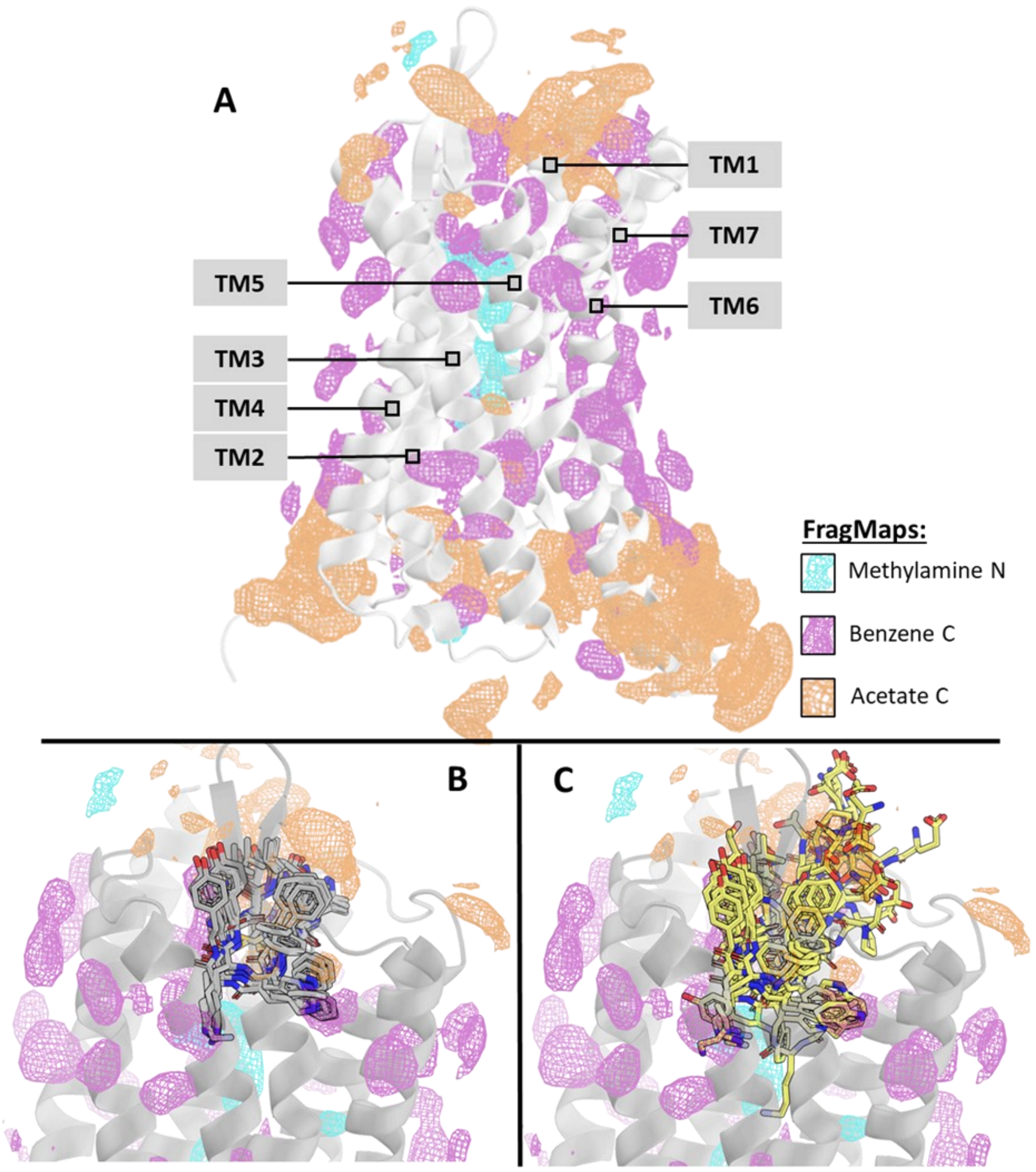


**Figure_S6**. (A) SILCS FragMaps of the active-state *h*UT model showing three main interaction regions: methylamine nitrogen FragMaps (blue) delineate a positively charged pathway through the receptor core; acetate carbon FragMaps (brown) are located at extracellular and intracellular sites; and benzene carbon FragMaps (purple) map the ligand-binding cavity and membrane-exposed surfaces. (B-C) Overlap of selected FragMaps with *h*UT and with the ten top-ranked poses of URP (B) or *h*UII (C).

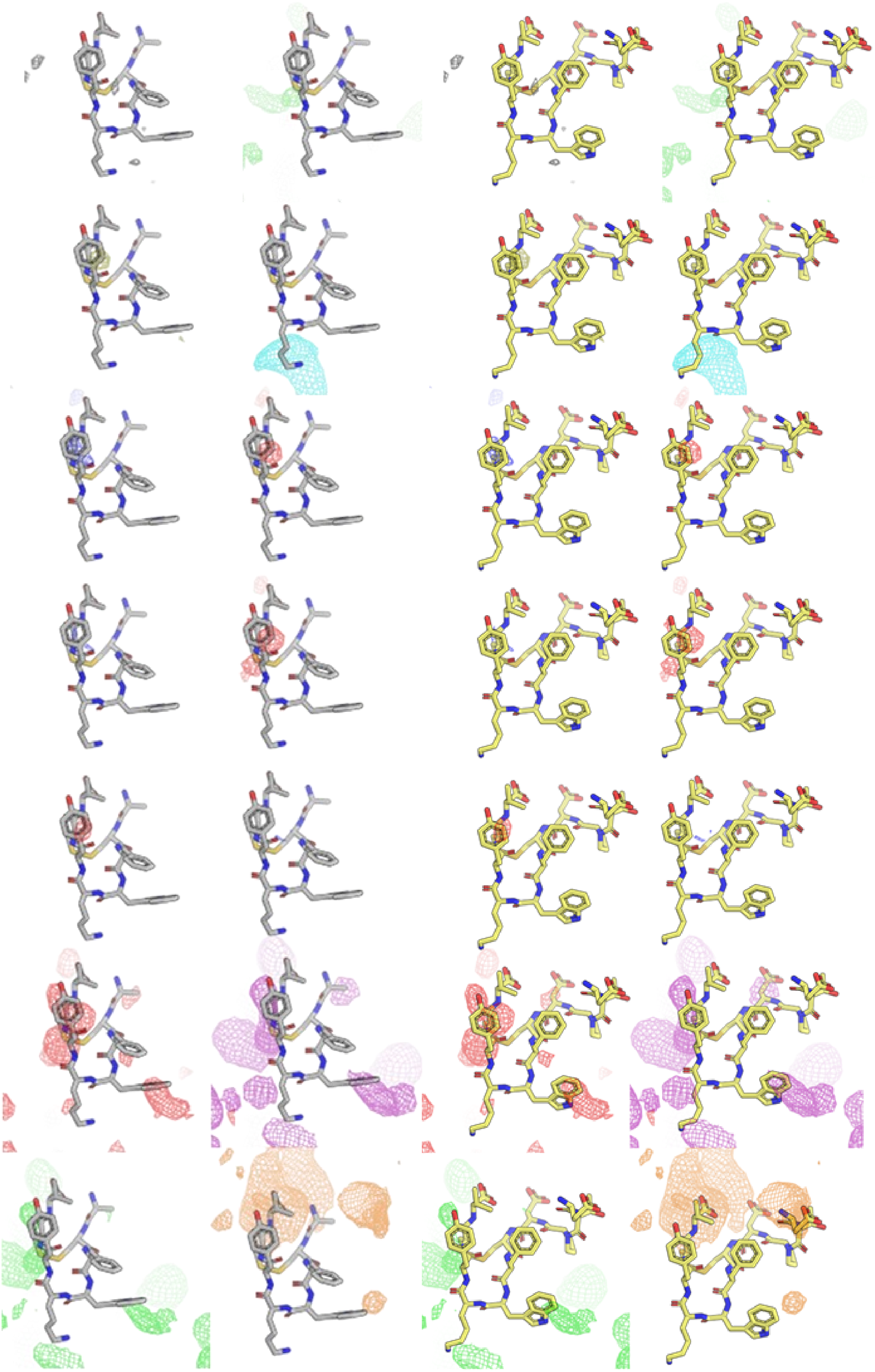

**Figure_S7**. Overlap of SILCS FragMaps with the highest-ranked docking pose of URP (left) and *h*UII (right). From left to right and top to bottom, the FragMaps shown correspond to: water oxygen; propane carbon; methanol oxygen; methylamine nitrogen; imidazole nitrogen (H-bond donor); imidazole nitrogen (H-bond acceptor); generic H-bond donors (formamide + imidazole); generic H-bond acceptors (formamide, dimethyl ether oxygen, imidazole nitrogen); formamide oxygen; formamide nitrogen; dimethyl ether oxygen; benzene carbons; generic apolar carbons (benzene carbons, propane carbon); and acetate carboxylate carbon.

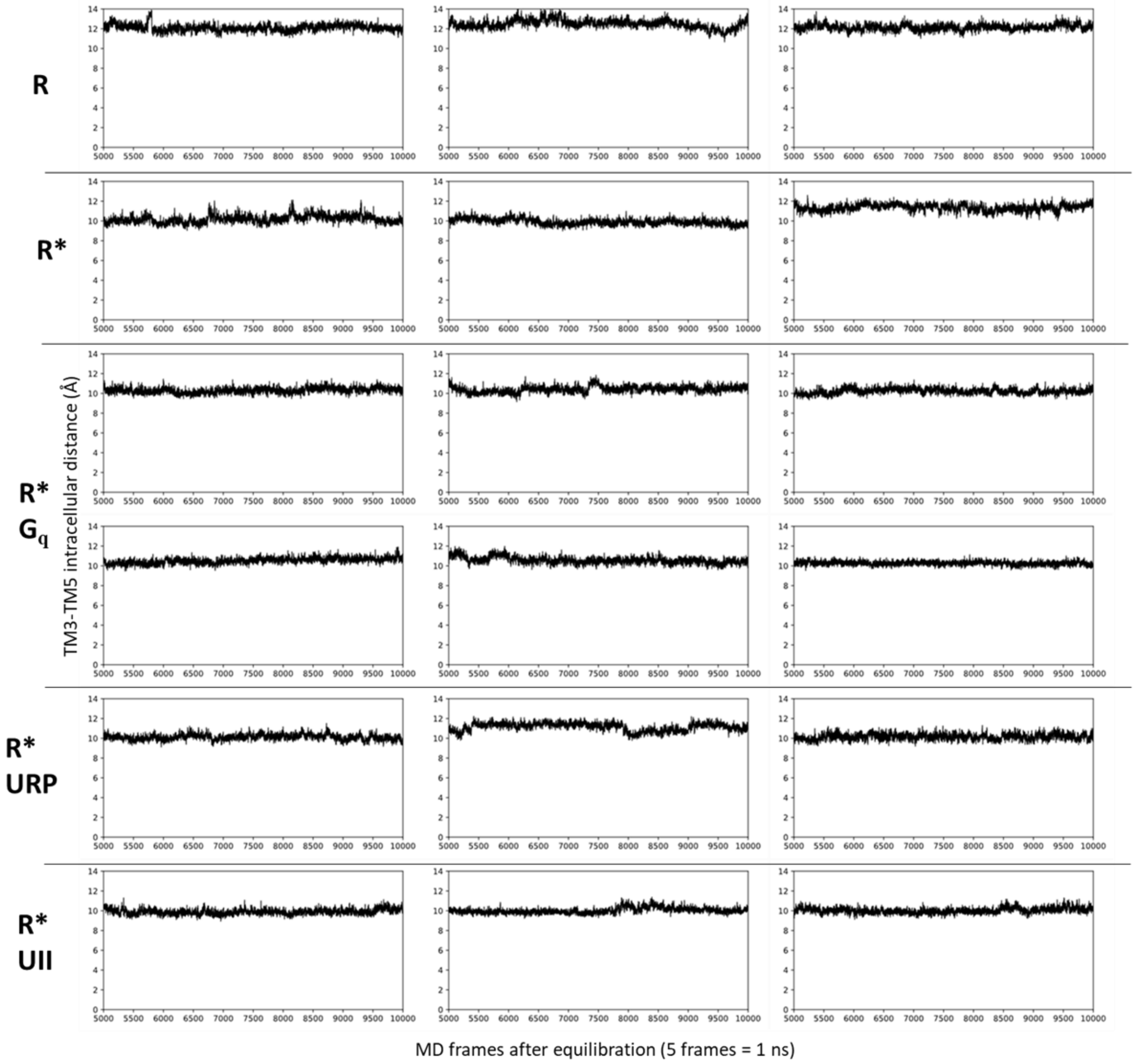


**Figure_S8**. TM3-TM5 time series in the final 1μs of each replicate simulation, each frame was saved every 200 ps, thus 10,000 frames correspond to the full 2000 ns trajectories. Inter-TM distances were measured as described in the *Methods* section.

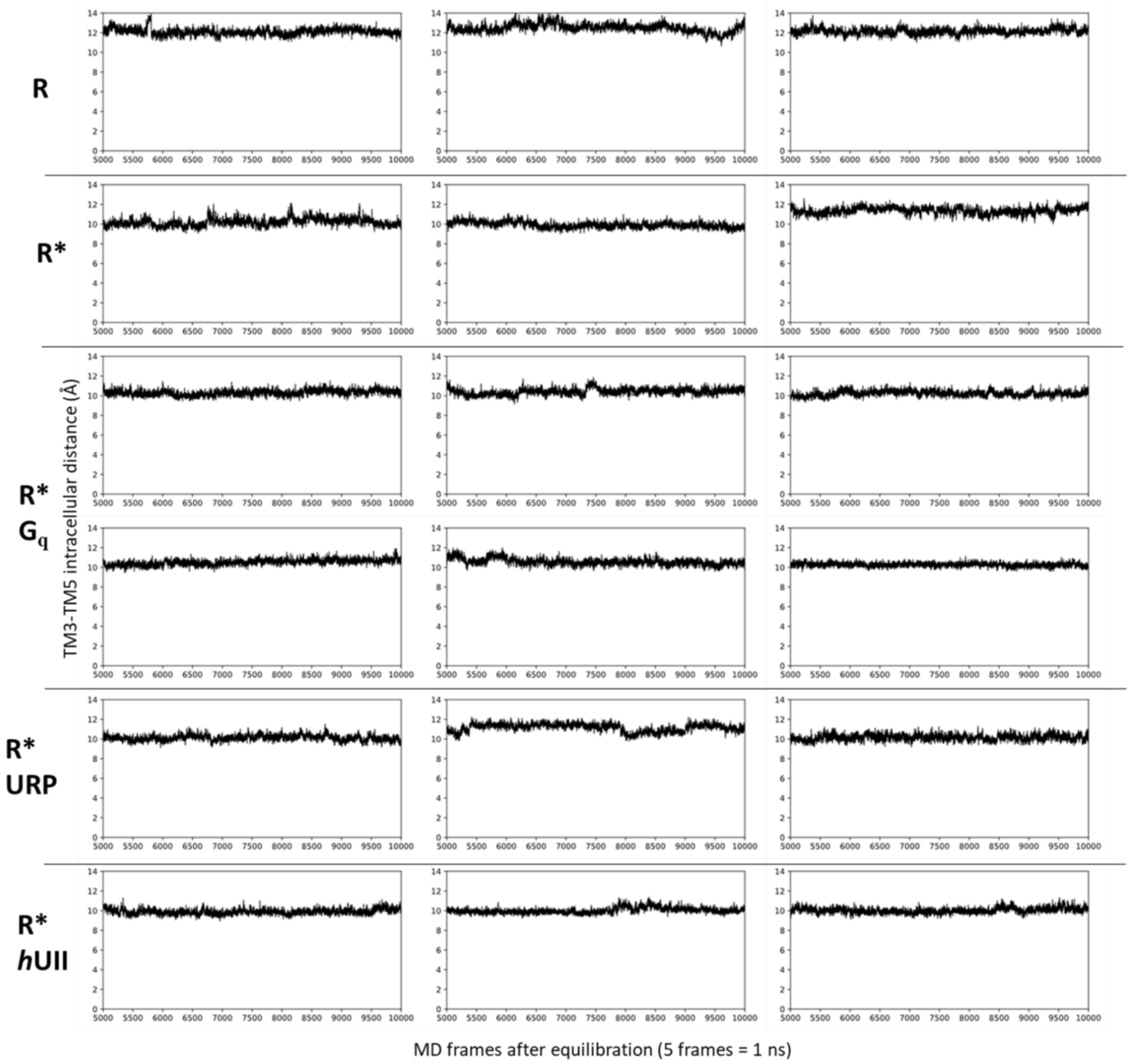


**Figure_S9**. TM3-TM6 time series in the final 1μs of each replicate simulation, each frame was saved every 200 ps, thus 10,000 frames correspond to the full 2000 ns trajectories. Inter-TM distances were measured as described in the *Methods* section.

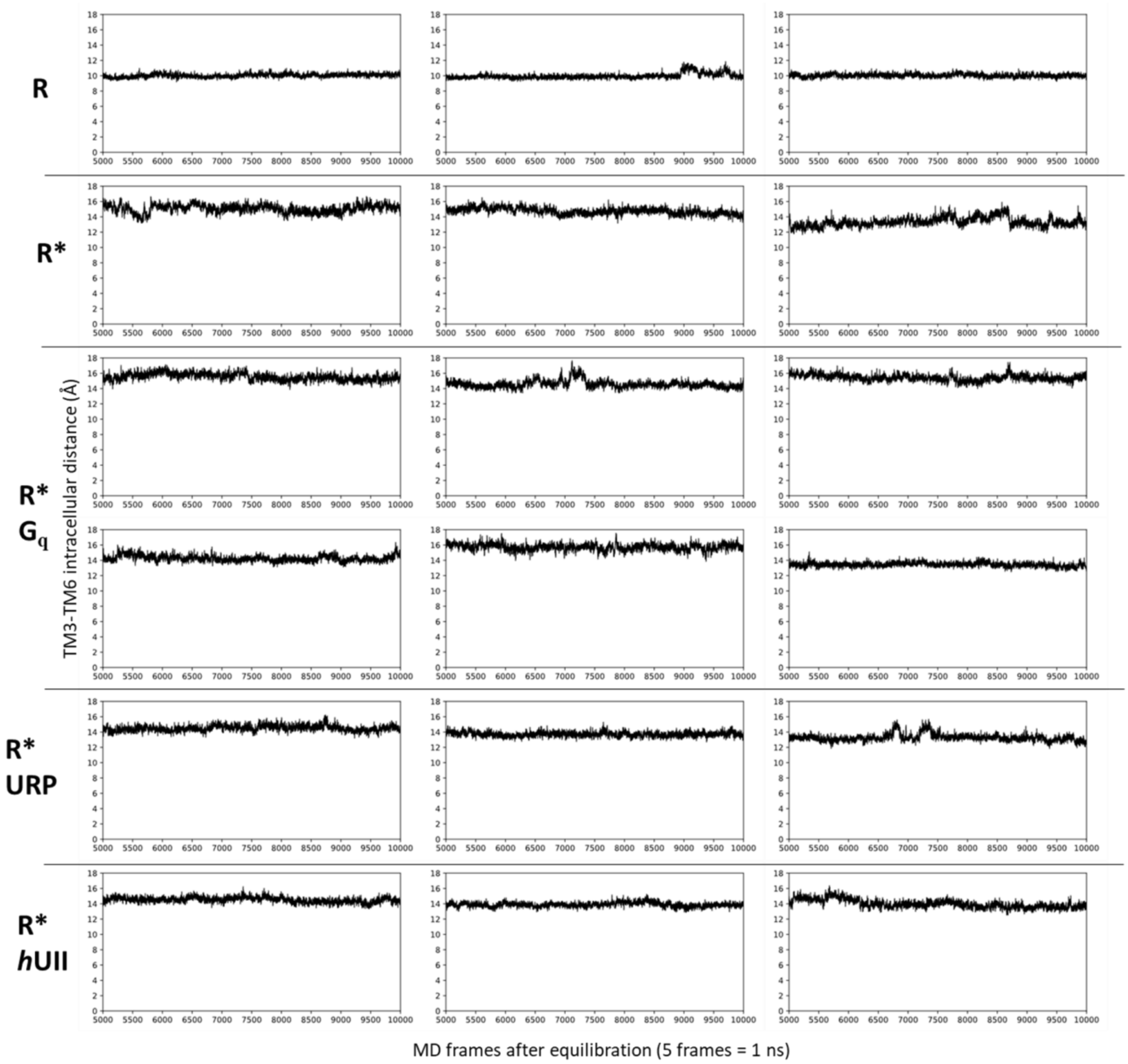


**Figure_S10**. TM3-TM7 time series in the final 1μs of each replicate simulation, each frame was saved every 200 ps, thus 10,000 frames correspond to the full 2000 ns trajectories. Inter-TM distances were measured as described in the *Methods* section.

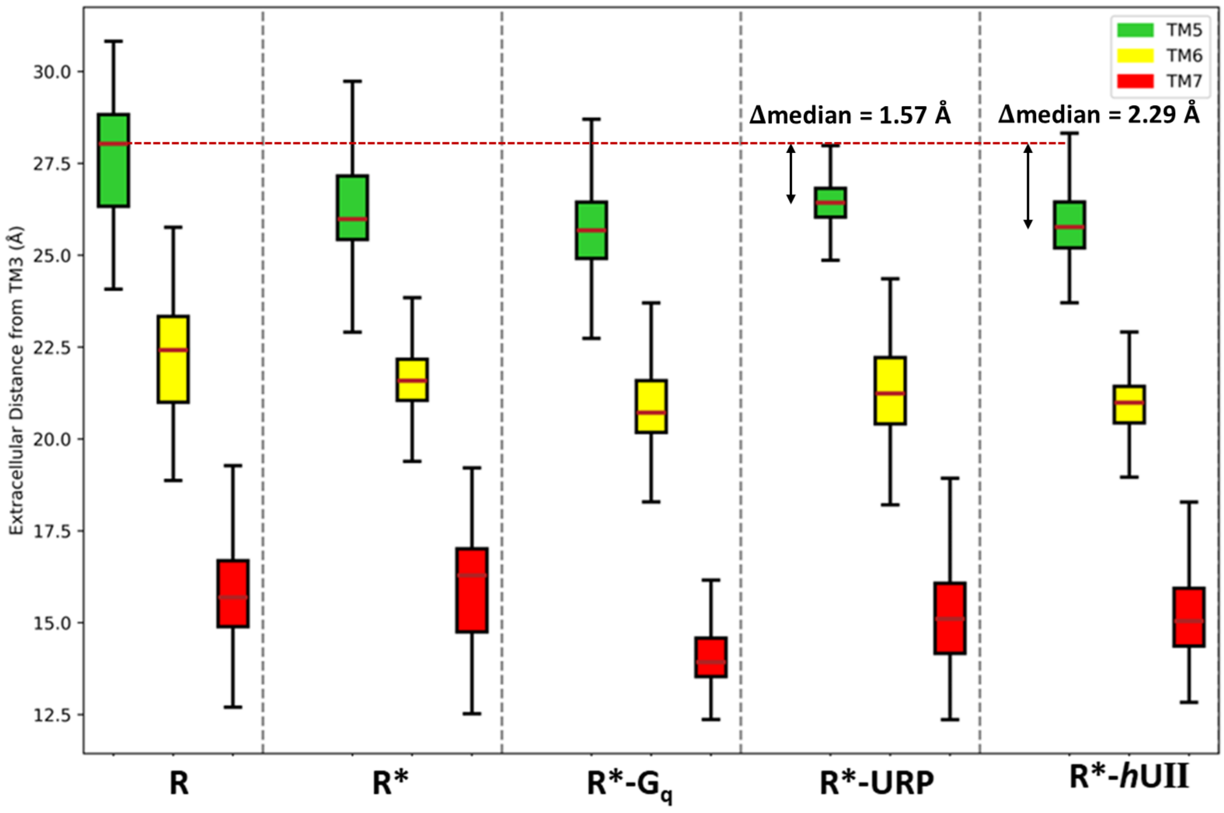


**Figure_S11.** Assessment of **extracellular** *h*UT dynamic TM-tilts are measured, the distribution of distances between the extracellular sections of TM5-7 to TM2 (see section *Methods*; TM2 is the most stable TM in the extracellular side) are plotted as box and whisker plots, the red full-line within the boxes indicates the median distance value, the whiskers indicate the maximum and minimum distances measured in the simulations. The red dotted line reports the median of the inactive state extracellular TM5 tilt to highlight its inward tilt upon activation and its ligand-dependent constraining by URP and *h*UII.

**Figures annotated for complete accessibility.** In order of appearance, these correspond to: figure 2, 4. In figure 2A, there are two kinds of FragMaps which are indicated by arrows, these are methylamine nitrogen and benzene carbon occupied volumes. In figures 4-7 the interaction objects created in PyMOL are straight translucent tubes that had to be color annotated to encode interaction subtypes due to the impossibility of shape-coding the interaction types. For each interaction its subtype is indicated by an arrow and text, except for hydrogen bonds which are encoded by shade, darker tubes indicate sidechain-to-sidechain hydrogen bonds, lighter tubes indicate sidechain-to-backbone and backbone-to-backbone hydrogen bonds.

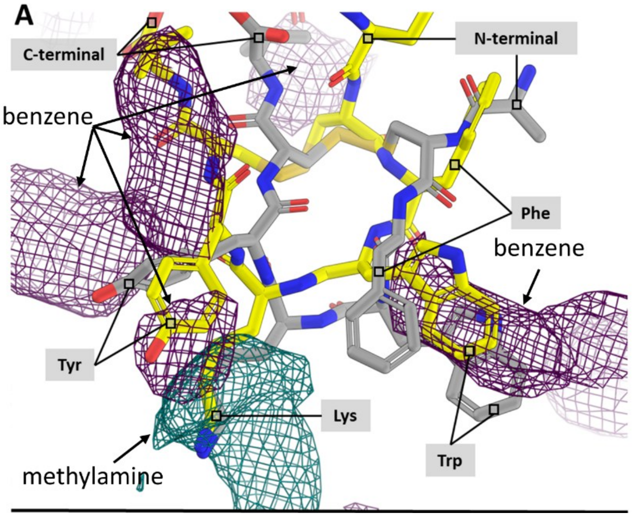


Color independent figure_2

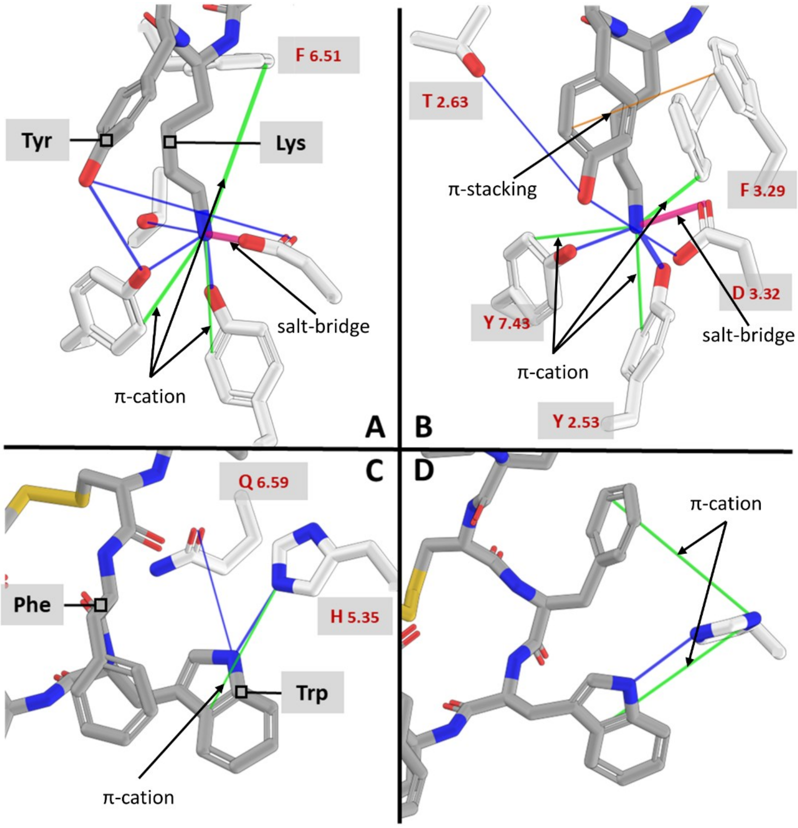


Color independent figure_4

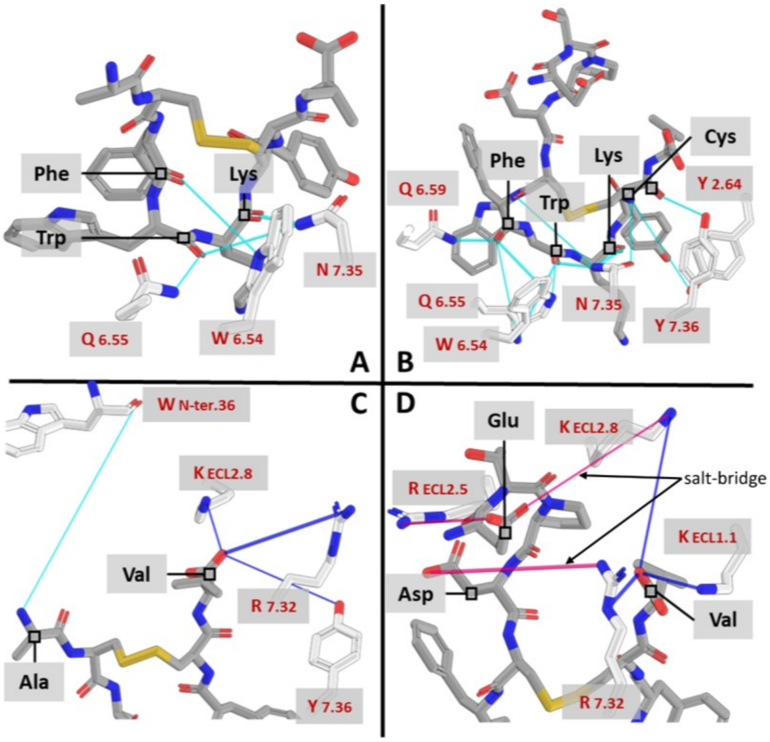


Color independent figure_5

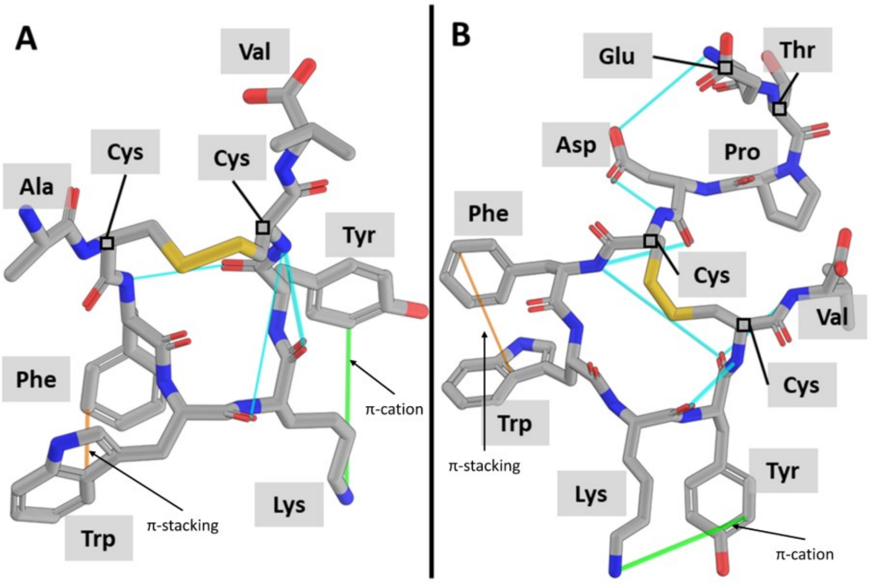


Color independent figure_6
